# A novel conserved protein associates with the IFT-A complex to mediate nuclear translocation of β-catenin in Wg/Wnt-signaling

**DOI:** 10.1101/2025.06.21.660855

**Authors:** Linh T. Vuong, Marek Mlodzik

## Abstract

Wingless (Wg)/Wnt signaling is critical throughout development and tissue homeostasis and associated with many disease states. Canonical Wg/Wnt-signaling is mediated by β-catenin (Arm in *Drosophila*) with IFT-A/Kinesin-2 complex promoting nuclear translocation of β-catenin/Arm upon pathway activation. Although IFT-A is essential for nuclear translocation of β-catenin/Arm, existing data suggests additional proteins are involved in this process. Here, we have identified a novel, evolutionarily conserved protein, Pasovec (Psv), as a critical component required for nuclear β-catenin/Arm localization. We demonstrate that Psv functionally interacts with the IFT-A/Kinesin2 complex and physically associates with IFT140, a core component of IFT-A. The Psv-IFT140 interaction is independent of Wg/Wnt-signaling activation. Importantly, Psv contains a nuclear localization sequence (NLS), which is critical for its own nuclear localization and that of β-catenin/Arm upon Wg/Wnt-signaling activation. Psv with a mutated NLS can act as an inhibitor of Wg/Wnt-signaling. Altogether, this study describes a new factor required for Wg/Wnt-signaling, with its mutant phenotypes resembling *wg* and *β-catenin/arm* mutants, that functions during the nuclear translocation process of β-catenin/Arm.

## Introduction

The canonical Wnt/Wingless (Wg)-signaling pathway is highly conserved across metazoans and required in the regulation of many processes, ranging from embryonic development and organogenesis, including tissue growth, cell fate specification, survival, migration, to stem cell maintenance and regeneration (rev in ^1–5^). Importantly, Wnt-signaling is also critically linked to several diseases, including developmental abnormalities, cilia associated defects, and initiation and progression of various types of cancers (rev in ^6–9^). In the absence of the Wnt signal, β-catenin/Arm is phosphorylated by the so-called “destruction complex” (DC) and thereby targeted for proteasomal degradation. The DC is composed of Axin, APC (Adenomatous Polyposis Coli), and the kinases GSK3β and CK1α (rev. in ^1, 3, 5, 10^). In canonical Wnt/Wg-signaling, Wnt binds to the Frizzled-LRP5/6 receptor/co-receptor pair leading to the disassembly of the DC, upon which cytoplasmic β-catenin (Armadillo/Arm in *Drosophila* ^11^) is released and therefore stable (e.g. rev. in ^3–5, 12^). Nuclear translocation of β-catenin/Arm is then the rate-limiting step in Wnt signal transduction for transcriptional target gene activation, with β-catenin/Arm being the key co-activator of the TCF/LEF transcription factor family (rev. in ^1, 3, 13^).

While the membrane proximal Wnt-signaling complexes, including receptor complexes, resulting in pathway activation and the composition and function of the DC are reasonably well understood (rev in ^1, 5, 10, 12^), understanding of nuclear translocation of β-catenin remains a work in progress ^14,15^. Recent work has identified a critical role for IFT-A (Intraflagellar Transport-A complex ^16, 17^) and Kinesin2 in the nuclear translocation of β-catenin/Arm ^18–20^. It has been established that a IFT-A/Kinesin2 complex binds to β-cat/Arm as it is released from the destruction complex, forming a transport complex that is required for nuclear translocation of β-cat/Arm ^18, 20^. IFT-A is generally known for its essential role during ciliogenesis and ciliary retrograde transport ^16, 17^. In Wnt-signaling IFT-A acts in the cytoplasm, and in particular it binds an N-terminal region of β-catenin/Arm (residues 34-87 in Arm, equivalent to the β-cat^24–79^ peptide in mammals) via its subunit IFT140 ^21^. IFT-A also binds to Kap3 connecting it to Kinesin 2/Kif3a ^18^. The resulting complex associates with cytoplasmic microtubules (MTs) and promotes nuclear translocation of β-catenin/Arm upon Wnt/Wg-pathway activation^18, 20^. Loss-of-function of the IFT-A complex components, Kap3, or Kinesin 2 results in impaired nuclear translocation of β-catenin *in vivo* in *Drosophila*, as well as in mouse embryonic fibroblasts and human cell lines ^18, 19, 21^. Importantly, the binding of β-catenin to IFT140 within the IFT-A is critical for nuclear translocation. Expression of the respective N-terminal β-catenin peptide interferes with IFT-A interaction with endogenous β-catenin by competing for/blocking its binding site, thus acting like an interference tool and suppressing Wg/Wnt-signaling *in vivo* during development and in cancer cell lines ^21^.

The surprising observation that the stable β-catenin/Arm mutant isoform, ArmS10 ^22^, which has this interaction region deleted, is still capable of localizing to the nucleus ^18^ and activating Wg-target gene expression, suggested that there are likely other binding proteins to β-catenin required for its nuclear translocation, either as cytoplasmic retention anchors, competing with IFT140 (withn the IFT-A) to inhibit the process, or additional positively-acting β-cat/Arm interacting factors that promote its nuclear translocation in parallel to or together with IFT-A. This hypothesis is further supported by the fact that ArmS10 nuclear localization is not affected by reduced IFT140 levels^18^. To identify additional potential factors required in this process, we searched the literature for proteins implicated in nuclear β-catenin/Arm localization and/or its nuclear function. This identified a gene called Twa1 (for Two hybrid-associated protein 1 with RanBPM)^23^, as a potential candidate, with Lu et al (2017) suggesting that it is required for nuclear retention of β-catenin in cancer cell contexts (^23^; see below for more details).

*Drosophila* wing development serves as a paradigm for Wnt/Wg-signaling dissection and analyses. In larval wing imaginal discs, Wg is expressed as a two-cell stripe at the dorso-ventral (D/V) compartment boundary of the wing pouch, where it acts as a morphogen and activates target genes in a concentration-dependent manner ^24–27^. Wg protein is detected in a gradient up to several cells away from its expression source^26, 27^, leading to a stereotypic pattern of gene expression governing wing development with its highest levels specifying wing margin cells ^28, 29^. Wg signaling targets in cells adjacent to the D/V boundary include *senseless (sens)*, a gene responsible for specification of wing margin sensory bristles ^30^, while lower Wg threshold target genes include *Distalless* (*Dll*), required for wing pouch growth and expressed in a graded fashion, decreasing away from the Wg expression stripe^28, 29^. Use of the wing imaginal disc patterning has thus served as a tremendous resource to dissect the Wnt/Wg signaling pathway and associated cell biological features^2^. In addition to the wing, *Drosophila* embryogenesis has served as an excellent tool in Wnt-pathway assembly, including the identification of *arm/β-catenin*, as a key component of Wg signaling (e.g. ^22, 31^).

Here we identify a conserved gene, *Drosophila CG18467* (mentioned as Twa1 or Gid8 [glucose-induced degradation protein homolog 8] in mammalian cells ^23, 32, 33^), which cooperates with the IFT-A/Kinesin2 complex in β-catenin/Arm nuclear translocation. Based on the embryonic loss-of-function phenotype of *CG18467*, which closely resembles that of *armadillo* (*arm*) mutants, we have named the gene *pasovec* (*psv*), the Czech word for armadillo. Our data suggest that *psv* is required for Wg/Wnt signaling in all tissues tested and critical for nuclear translocation of β-catenin/Arm. Our results are consistent with a model where Psv directly associates with β-catenin/Arm to promote its nuclear localization, at least in part in a cooperative manner with the IFT-A complex. Strikingly, Psv contains a nuclear localization signal (NLS), which we show to be critical for its role in Wg/Wnt signaling and β-catenin/Arm nuclear localization, both in *Drosophila* and in human (with the human orthologue Gid8 performing the equivalent function). Our work described here not only identifies a new conserved factor in Wg/Wnt-signaling but also provides new insight into an emerging mechanistic model for nuclear transport of β-catenin/Arm upon Wnt-signaling activation.

## Results

### *CG18467*/*pasovec* knock-down resembles Wg-signaling defects

As mentioned above, the observation that the stable ArmS10 mutant isoform ^22^ is capable of translocating into the nucleus led us to screen for potential additional factors in the process of nuclear β-cat/Arm translocation. A literature search identified a gene called Twa1 (for Two hybrid-associated protein 1 with RanBPM [Ran Binding Protein M]) ^23^, as a candidate. Twa1, *CG18467* in the *Drosophila* genome, is highly conserved across animal species, exhibiting 68% sequence identity between the *Drosophila* and human proteins (Suppl. Fig. S1A), and contains a C-terminal domain proposed to interact with RanBPM ^23^ (underlined in Fig. S1A). We tested the equivalent *Drosophila* gene, *CG18467*, for a potential role in Wg-signaling. As a first approach, we employed the UAS/Gal4 system ^34^ for RNAi-mediated gene knockdown at the wing margin, using *C96-Gal4*, which serves as a rigorous model for Wg-signaling dissection. *C96-Gal4*, drives expression in cells at and surrounding the dorsal-ventral (D/V) boundary in wing imaginal discs (^35^; also Suppl Fig. S1B). RNAi-based knockdown of *CG18467* (referred to as *pasovec/psv*, see below for explanation) with three independent RNAi-lines using *C96-Gal4* resulted in loss of wing margin cells and associated notching phenotypes, consistent with impaired Wg pathway activity (Fig. 1A-C, with increased temperature and knock-down causing stronger defects; also Suppl. Fig. S1C-D).

**Figure 1.**
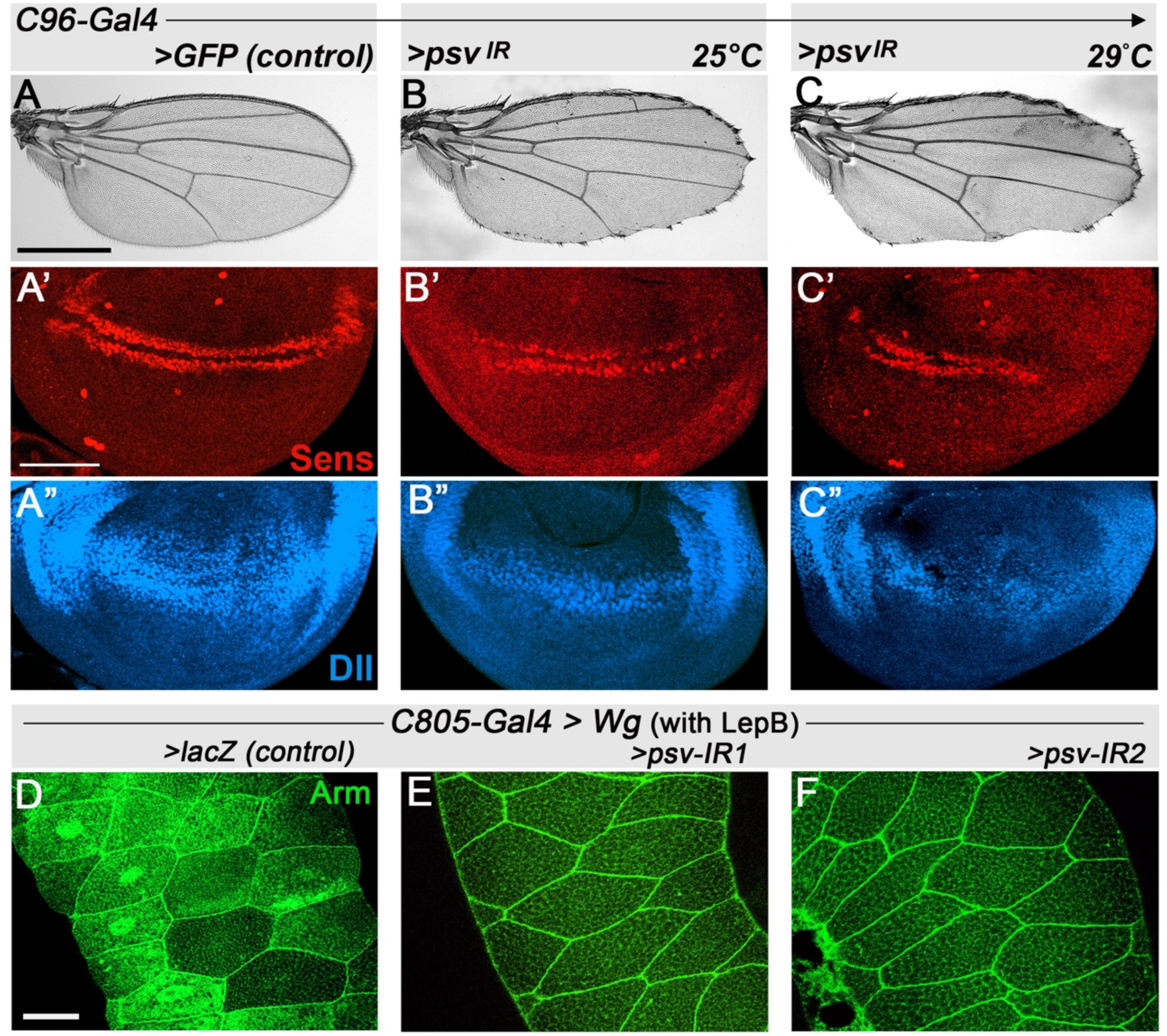
*psv* is required for wing margin development and nuclear ý-cat/Arm localization. (A-C) Wing development effects of *psv* RNAi knockdown with *C96-Gal4* at temperatues as indicated. (A-A”) Control wing and wing discs with *C96>GFP* showing normal adult wing (A), and wing disc with Sens expressing cells (red, A’) and Dll expression (blue, A”). *psv* RNAi (*C96>psv^IR^*) reveals partial loss of wing margin (B,C). (B’-C’) *C96>psv^IR^* causes reduction/loss of Sens expressing cells (B’), further reduced at 29°C (C’). (B”-C”) Expression of Dll near DV boundary (shown in blue) is reduced in *C96>psv^IR^* (B”, C”). (D-F) Wg-signaling induced ý-cat/Arm nuclear translocation assay in salivary glands (SGs). Wg was expressed via *C805-Gal4*; note mosaic expression, common to all SG drivers (see text for reference) treated with LepB (nuclear export inhibitor) to enhance nuclear Arm/β-catenin retention. SGs were stained for endogenous Arm (green). (D) Wg expression (*>Wg, >lacZ*: positive control): note increased cytoplasmic Arm/β-catenin and its nuclear localization (uneven Arm/β-cat levels due to mosaic Gal4 driver expression). (E-F) Knockdown of *psv* (two independent RNAi lines) in the *C805>Wg, >psvRNAi* background. Note that nuclear translocation of endogenous Arm/β-catenin is blocked (junctional Arm serves as internal control, marking cellular outlines). Scale bar represents 500μm in adult wings (A-C), 50 μm in imaginal discs (A’-C”), and 30 μm in salivary glands (D-F).

To further investigate the role of *CG18467/psv* in wing development, we examined its requirement for the expression of key Wg/Wnt-signaling target genes *senseless* (Sens, ^30^) and *Distal-less* (*Dll*, ^36^). RNAi knockdown of *CG18467/psv* (using *C96-Gal4*) led to a marked reduction in the number of Sens expressing cells and Dll expression at the D/V-boundary (Fig.1A’-C”; note further reduction with increased temperature, causing a stronger knockdown). There was no detectable induction of Caspase 3 in response to *CG18467/psv* RNAi near the D/V boundary region (Suppl. Fig. S1F-F”). These results suggest that *CG18467/psv* is specifically required for Wg/Wnt-signaling and Wg target gene activation and associated wing patterning.

As the premise of this study was to identify additional factors required in the nuclear translocation process of β-cat/Arm, we next assessed *CG18467/psv* in the salivary gland (SG) assay, which due to the large size of SG cells and nuclei facilitates detection of nuclear β-cat/Arm upon Wg induction^18^. Strikingly, Wg-induced β-cat/Arm translocation was eliminated by RNAi-mediated knockdown of *CG18467/psv* (Fig. 1D-F). Based on the above observations, along with the embryonic loss-of-function (LOF) phenotype of *CG18467/psv* (see below), we have named the gene *pasovec* (*psv*, the Czech word for armadillo).

### *pasovec* (*psv*) LOF mutant phenotypes display Wg-signaling defects

To confirm and further define the role of *psv* as a Wg-signaling pathway factor, we have generated LOF alleles of the gene, using CRISPR/Cas9 methodology (see Methods), thus generating two independent LOF alleles, designated as *psv^null^* and *psv^λ1C^* (see Suppl. Figure S2A for allele information, both alleles show inditiguishable phenotypes). Analyses of *psv* mutant tissue in wing imaginal discs revealed that expression of the Wg-target genes Sens and Dll is largely eliminated in *psv* mutant cells (Fig. 2A-E’; note that Dlg as cellular junctional outline is not affected, indicating that *psv* does not affect epithelial cell architecture). These data were consistent with the RNAi experiments for *psv* shown above (Fig. 1).

**Figure 2.**
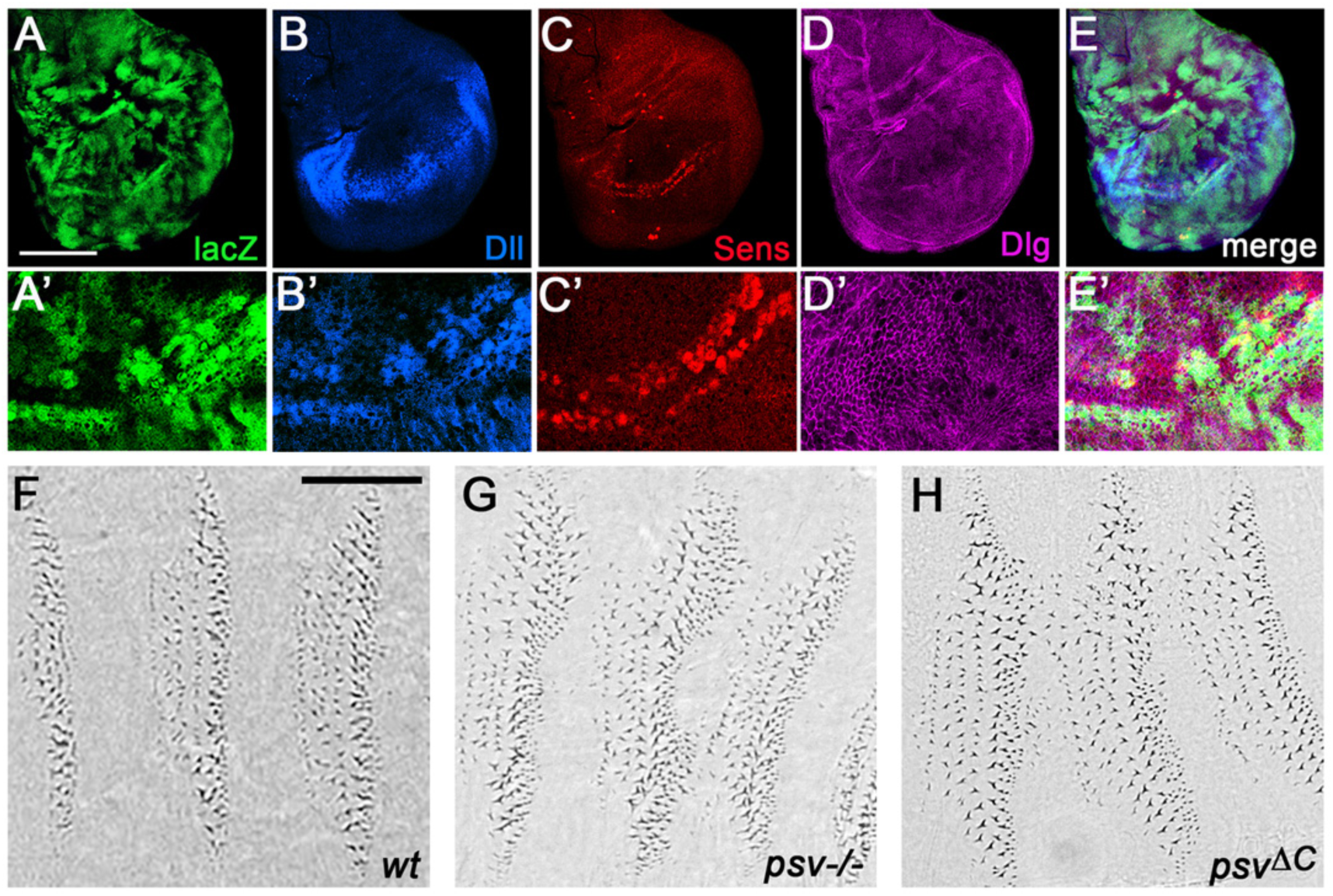
*psv* LOF mutant analyses in wing discs and embryo patterning. (A-E’). *psv^-/-^* (null allele) mutant clones in wing discs, marked by absence of *lacZ* expression (anti-β-Gal, green in A-A’), anti-Sens (red, B-B’), anti-Dll (blue, C-C’) and anti-Dlg (magenta, D-D’). (A’-E’) show high magnification of wing margin region. Note Sens and Dll expression is largely lost in *psv^-/-^* clones, with Dlg (control) appearing normal at cell membranes (D, D’). (E, E’) are merged images of all channels. (F-H) *psv* is required for embryo patterning. (F) High magnification view of ventral denticle belts in wild-type (F; note normal anterior-posterior denticle pattern). (G-H) Both *psv* mutant alleles, *psv^-/-^* (null allele, G) and *psv^λ1C^* (H), display defective patterning of denticles belts, reflecting a segment polarity phenotype comparable to *wg* and *arm* mutants (see main text for references), with ectopic denticles forming in segmental areas that are normally present with naked cuticle. Scale bar represents 50μm (A-E) and 100*μm* (A’-E’) in wing imaginal discs, and 80μm in embryos (F-H).

In addition to wing patterning, Wg signaling is essential in many other contexts, and well known for establishing segmentation and segment polarity during embryogenesis, giving the *armadillo* gene its name (e.g. rev. in^37, 38^). In this context, loss of *arm* or other Wg signaling components disrupts the anterior-posterior segmental patterning and associated denticle specification on the ventral epidermis of embryos and larvae^39^, rev. in^36^). Knockdown of *psv* using *daughterless-Gal4* (*da-Gal4*, driving expression in all embryonic cells) resulted in a consistent segment polarity phenotype, characterized by ectopic denticles and partial loss of naked cuticle (Suppl. Fig. S2B-B’, showing segments A4-A6), very similar to the phenotype observed in *da-Gal4, UAS-arm^RNAi^* embryos (Suppl. Fig S2B”). Confirming this phenotype and requirement of *psv* for Wg-signaling in embryonic patterning, both homozygous *psv* LOF alleles, *psv^null^* and *psv^λ1C^*, disrupted denticle patterning, causing segment polarity type segmentation defects (Fig. 2G-H, compare to Fig. 2F for wild-type anterior-posterior denticle pattern), which were very similar to *arm* LOF alleles that specifically affect Wg-signaling ^11^. This phenotype gave *pasovec* (*psv*) its name.

Taken together with the requirements in wing development and the salivary gland assay, these data indicate that *psv* plays a general role in Wg/Wnt signaling (in all tissues tested), likely by affecting the nuclear translocation of Arm/β-cat.

### *psv* functionally interacts with the IFT-A/Kinesin2 complex

Since a *psv* knockdown disrupts the nuclear translocation of Arm/β-cat (Fig. 1D-F), we next tested whether it acts together with the IFT-A/Kinesin2 complex. To assess this, we examined whether *psv* functionally interacts with components of this complex, using interference and overexpression in the wing margin specification assay. Very similar to *psv^RNAi^* knockdown (Fig. 3A-B; via RNAi expressed under *C96-Gal4* control), mild knockdown of IFT140, the Arm/β-cat interacting component of IFT-A, or Klp64D/Kinesin2, resulted in a partial loss of the wing margin (Fig. 3C and 3F, respectively). Reducing IFT140 or Klp64D levels in the *C96-Gal4, UAS-psv^RNAi^*background enhanced the respective phenotypes, leading to an increase in loss of wing margin cells and wing tissue (Fig. 3D and G, compare to 3B and 3C for example). Importantly, over-expression of IFT140 or Klp64D (in the *C96-Gal4 UAS-psv^RNAi^* background) largely suppressed the margin defects induced by *psv^RNAi^* (Fig. 3E and H; note that *C96*-driven overexpression of IFT140 or Klp64D alone did not cause any detectable defects in adults or Sens and Dll expression in wing discs, Suppl. Fig. S3).

**Figure 3.**
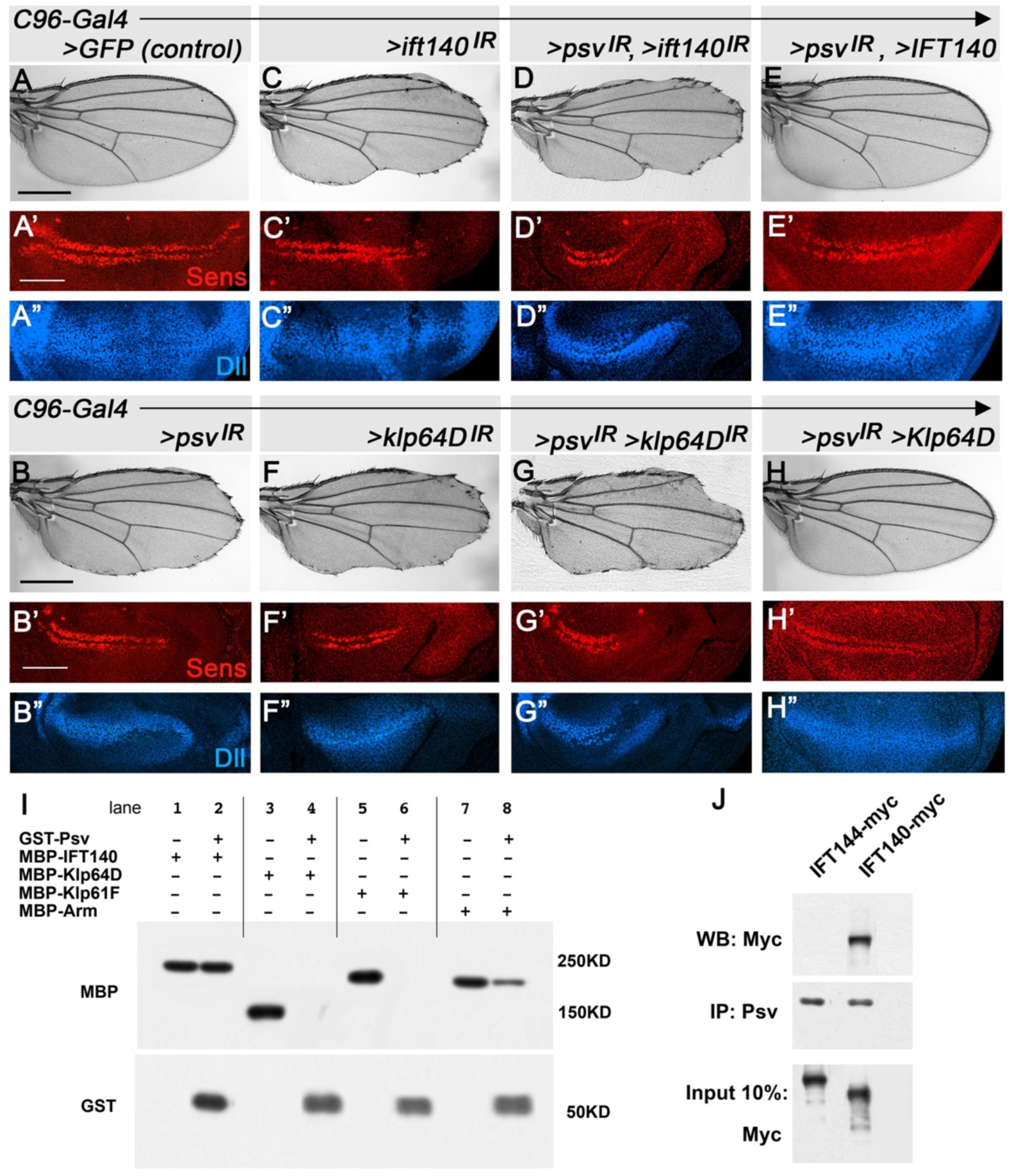
*psv* interacts functionally with IFT-A/Kinesin2 complex. (A-H”) Functional interactions during wing development. (A-A”) Control (*UAS-GFP*: *C96>GFP*) adult wing (A) and wing disc with Sens (red, A’) and Dll (blue, A”) expression near DV boundary. (B-B”) Equivalent *psv* knockdown (*C96>psv^IR^*) wing and wing disc, with partial loss of margin (B), reduction of Sens expressing cells (B’) and reduction in Dll expression (B”). (C-H”) Interactions between *psv* and *ift140* (for IFT-A) or *klp64D* (for Kinesin2). (C-C”) *C96>ift140^IR^* control and (F-F”) *C96>klp64D^IR^* control: partial loss of wing margin (C,F), reduction in Sens expressing cells (C’,F’) and Dll expression near DV boundary of wing discs (C”,F”), indistinguishable from >*psv^IR^* (cf. to B-B”). (D-D”) and (G-G”): *C96>psv^IR^, >ift140^IR^* (D-D”) or *C96>psv^IR^, >klp64D^IR^*(G-G”) double knockdowns; note increased wing margin defects, further reduction in Sens expressing cells (D’,G’) and Dll expression (D’,G”); cf. to (B-B”) and (C-C”). (E-E”) and (H-H”): Overexpression of either IFT140 (*C96>psv^IR^, >*IFT140) or Klp64D (*C96>psv^IR^, >Klp64D*) in *C96>psv^IR^* background, suppressed adult phenotypes (E,H) and restored Sens (E’,H’) and Dll (E”,H”) expression. Scale bar represents 500 μm in adult wings (A-H), and 50 μm in imaginal discs (A’-H”). (I-J) Psv physically associated with Arm and IFT140. (I) Direct binding of GST-Psv to IFT140 and Arm (all as MBP-fusions: input 10% shown in odd numbered lanes). GST-Psv pulled down IFT140 (lane 2) and Arm (lane 8), whereas full-length MBP-Klp64D and MBP-Klp61F were not pulled down (lanes 4 and 6). (J) *In vivo* co-immunoprecipitation (co-IP) assay of IFT144 (negative control) and IFT140 with Psv, from *nub-Gal4>IFT144myc* and *nub>IFT140myc* wing imaginal discs. Protein extracts from wing discs were IPed with anti-Psv (input 10% of wing disc lysates used in co-IP) and analyzed by blotting with anti-myc to IFT144 or IFT140.

Consistent with wing margin loss, knockdown of *psv* analyzed in wing discs resulted in reduction of Sens expressing cells and Dll expression along the D/V boundary, and within the wing pouch respectively (Fig. 3B’-B”, compare to wild-type control in Fig. 3A’-A”; see also Sens expression quantification in Suppl. Fig. S3). The reduction in Sens and Dll expression caused by *psv^RNAi^* was either enhanced by concomitant knockdown of Klp64D (Kinesin2) or IFT140 (Fig. 3D’-D” and 3G’-G”, respectively) or, importantly, rescued by their overexpression (Fig. 3E’-E” and 3H’-H”, respectively; compare to wild-type, Fig. 3A’-3”). Similar interactions were observed with Kap3 (Suppl. Fig. S3), the Kinesin2 adapter required in nuclear Arm/β-cat translocation^14, 18^. Taken together, these data suggested that Psv functions together with the IFT-A/Kinesin2 complex in Wg/Wnt-signaling and either of the components can be rate limiting.

### Psv directly associates with IFT140 and Arm/β-cat

To refine the mechanism of the interaction between Psv and the IFT-A/Kinesin2 complex, we tested whether Psv physically associates with components of this protein complex. Strikingly, GST-Psv displayed direct physical interaction with IFT-A, via IFT140 (Fig. 3I, lanes 1-2), and Arm/β-cat (Fig. 3I, lanes 7-8), as demonstrated by *in vitro* pull downs of the respective MBP fusions (Fig. 3I, odd lanes represent 10% input and even lanes the pulldowns). These interactions were specific, as Psv did not pull down Kinesin2 (Klp64D, *Drosophila* Kif3a; Fig. 3I, lanes 3-4) or a control kinesin (Klp61F; Fig. 3I, lanes 5-6). To confirm the interaction between Psv and IFT140 *in vivo* during wing patterning, we performed co-immunoprecipitation (co-IP) assays with *Drosophila* wing imaginal disc extracts expressing Psv-GFP and IFT140 (or IFT144 as negative control; under *nub-Gal4* expressed throughout the wing pouch region). Extracts from *nub>IFT140-myc* and *nub>IFT144-myc* wing discs were IPed with anti-Psv antibodies (see below and Methods) and probed with anti-myc (Fig. 3J). This *in vivo* experiment revealed that IFT140 readily co-IPed with Psv (Fig. 3J; in contrast, the control, IFT144, did not IP with Psv).

Taken together these results support the conclusion that Psv can directly interact with both IFT140/IFT-A and Arm/β-cat, and further suggest that Psv plays a role in canonical Wg/Wnt signaling and associated nuclear translocation of Arm/β-cat by participating in the IFT-A/Kinesin2 complex mediated process.

### Psv shows overlapping subcellular localization with the IFT-A complex *in vivo*

The genetic and physical interactions between Psv and the IFT-A/Kinesin2 complex indicated that these proteins might function together in the same complex or subcellular compartment. To address this *in vivo*, we generated a specific antibody against Psv, which recognizes endogenous Psv protein in both Western blots and *in vivo* in fixed tissues (Suppl. Fig. S4; see Methods for details). We examined the subcellular localization of Psv and co-stained with IFT140-myc or Klp64D-HA (Kinesin 2) in *Drosophila* wing imaginal discs and naïve salivary gland cells. In wing discs, Psv was detected in intracellular puncta in cells near the wing margin (Fig. 4A-B), where Wg-signaling is active (also Suppl. Fig. S4). Notably, these puncta co-stained quite extensively with IFT140-myc (Fig. 4A’, A’’’, for additional examples see Suppl. Fig. S4D), and also displayed partial overlap with Klp64D-HA staining (Fig. 4B-B’’’; Suppl. Fig. S4E). While the observed co-localization in wing imaginal disc cells is in a tissue that is exposed to Wg-signaling, we also wished to analyze this in ‘naïve’ cells that are not exposed to Wg. The salivary gland cells serve this purpose, as they only get exposed to Wg when it is induced by the Gal4-system. In ‘naïve’ salivary gland cells, in absence of Wg, endogenous Arm is only detected at adherens junctions (Fig. 4C” and 4D”). Strikingly, in the same cells, a partially overlapping localization of Psv and the IFT-A complex (as detected with IFT140-HA) is observed in cytoplasmic puncta (Fig. 4C-C’’’). Similarly, partial cytoplasmic co-localization to puncta was detected between Psv and Kinesin2 (Klp64D-HA; Fig. 4D-D”’; note that Klp64D-HA also associates with adherens junctions in salivary gland cells, unlike Psv and IFT-A; Fig. 4D’).

**Figure 4.**
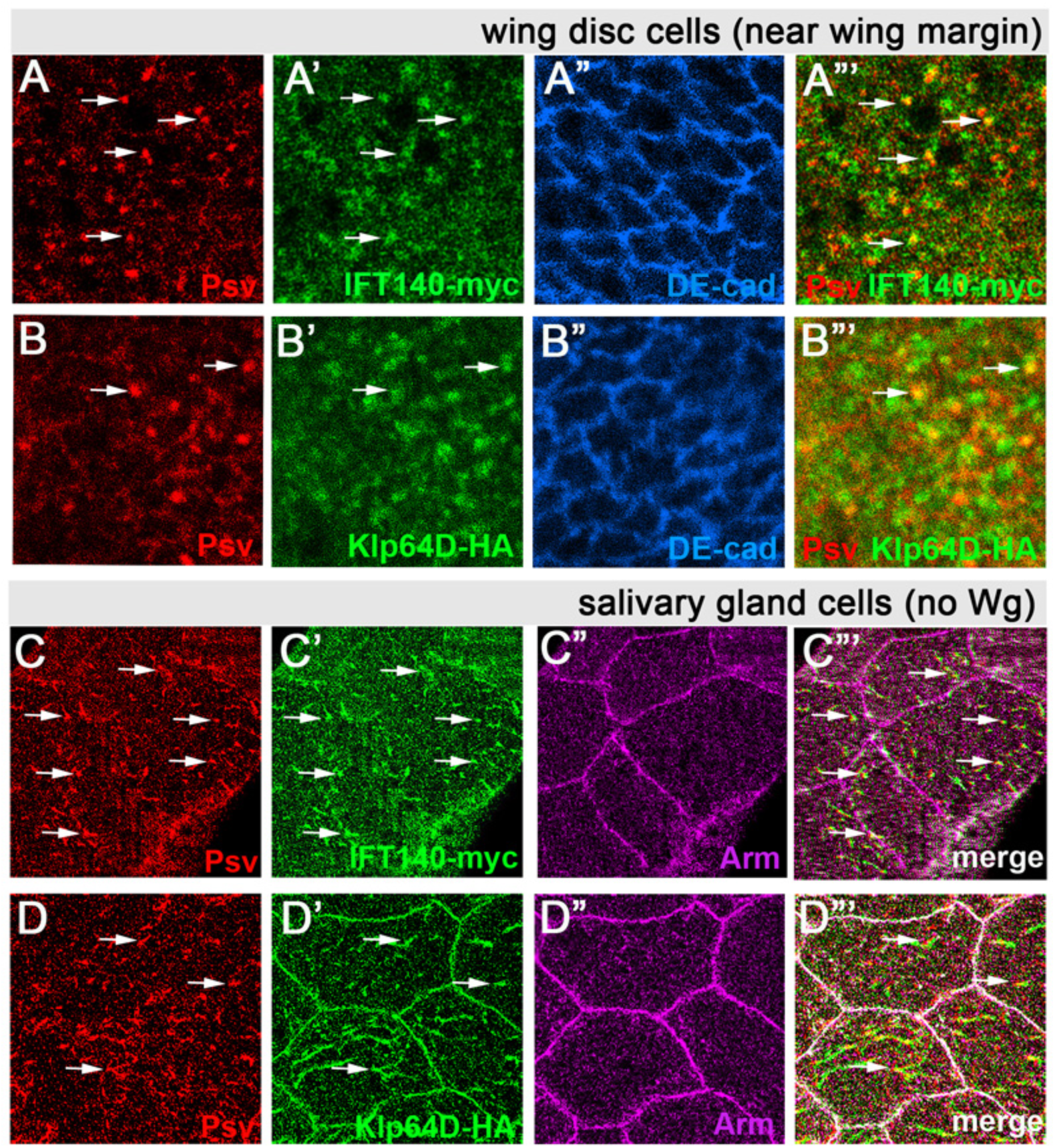
Psv and Kinesin2/IFT140 co-localize in wing discs and salivary gland cells. (A-B”’) Localization of Psv (red, anti-Psv) and IFT140-myc (myc, green A’ and A’’’) and Klp64D-HA (anti-HA, green in B’ and B’’’) in cells near the D/V boundary of wing discs. *C96>IFT140-myc, Klp64D-HA* wing disc were co-stained with anti-Psv. DE-cad (blue in A” and B”) serves as junctional marker revealing cellular outlines. Arrows mark examples of puncta staining for Psv overlapping with IFT140-myc (in A, A’, and A”’) and Klp64D-HA (in B, B’, and B”’). (A”’ and B”’) show merged images of green and red channels. (C-D’’’) Kinesin2/IFT140 complex and Psv co-localize in naïve salivary glands (SGs) independent of Wg signaling. (C-C”’) SGs stained for Psv (red), IFT140-myc (green) and endogenous Arm (magenta). Without Wg, Arm/β-catenin localizes to adherens junctions (AJs). Note that Psv and IFT140 partially co-localize, displaying punctate cytoplasmic staining (arrows: examples of co-stained puncta). (D-D’’’) SGs stained with Psv (red), Klp64D-HA (Kinesin2, green), and endogenous Arm. Note that Klp64D-HA localizes also to AJs, like Arm, and partially overlaps with Psv in cytoplasmic puncta (arrows: examples of co-stained puncta). See Fig S4 for quantification. Scale bars equal 10μm for wing imaginal discs (A-B’’’), and 30μm for salivary glands (C-D’’’).

Taken together, these data indicate that (i) Psv associates with the IFT-A complex (IFT140) in the cytoplasm in the presence or absence of active Wg-signaling, and (ii) Psv does not localize to AJs, and therefore does not associate with junctional Arm/β-cat in either cell type tested. These observations suggest that Psv forms complexes with IFT140 independently of Wg/Wnt signaling, and that it only associates with Arm/β-cat upon Wg/Wnt-signaling exposure (see below).

### The NLS of Psv is required for its role in Wg/Wnt-signaling *in vivo* and in cancer cells

The Psv protein contains a nuclear localization sequence (NLS) within its C-terminal RanBPM interacting domain, which is highly conserved from *Drosophila* to human (see Suppl. Fig S1 for sequence alignment; residues in green mark the NLS sequence). We asked whether this NLS is functional by testing whether it promotes nuclear localization of Psv. To this end, we mutated the respective arginine residues within the RRAFDRPE motif to alanine (AAAFDRPE, see Fig. 5A) and compared the wild-type, Psv-GFP, and mutant protein, PsvNLS*-GFP, in salivary glands or in S2 cells. Expression of wild-type GFP-tagged Psv in salivary gland cells (*C805-Gal4, UAS-Psv-GFP*) displayed a predominantly nuclear localization (Fig. 5B). In contrast, Psv with a mutated NLS (PsvNLS*-GFP) in the same assay showed strong cytoplasmic localization and even nuclear exclusion (Fig. 5C). Similarly, the same proteins transfected into S2 cells displayed either enriched nuclear localization (Psv-GFP) or nuclear exclusion (PsvNLS*-GFP; see Suppl. Fig. S5A-B), establishing that the NLS is functional and likely required.

**Figure 5.**
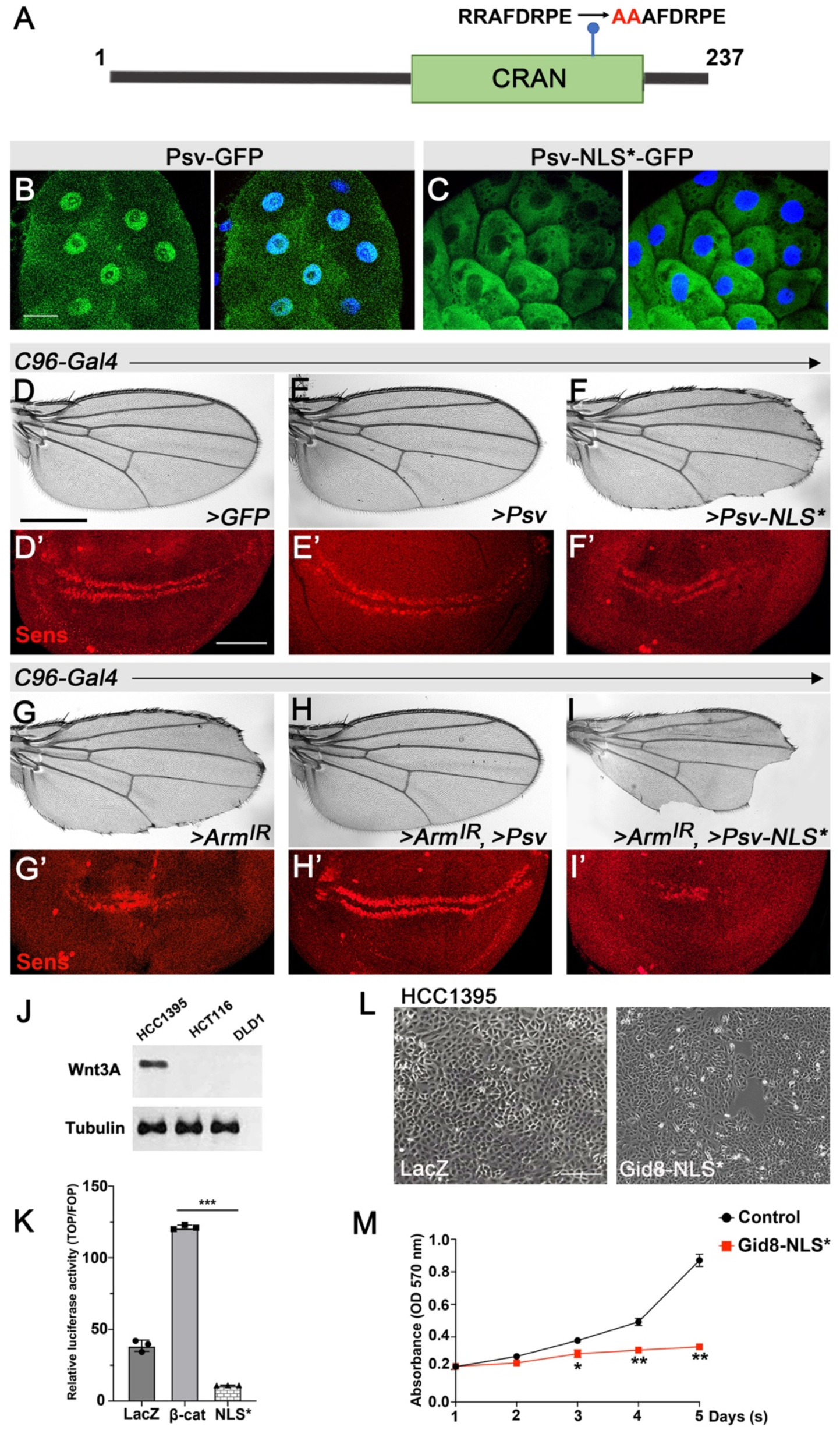
Nuclear localization sequence (NLS) of Psv is required for its function in Wg/Wnt-signaling. (A) Sequence of NLS in the C-terminal region of Psv, CRAN (C-term RanBPM binding domain) (see Fig. S1 for sequence alignment) and its mutant form, NLS*, with substitution of Arginines 144, 145 to Alanine. (B-C) Subcellular localization of overexpressed Psv-GFP (B) and PsvNLS*-GFP (C) in SG cells (green, DAPI nuclear staining in blue). Overexpressed Psv-GFP is predominantly detected in nuclei (note that this is different from wild-type Psv, and likely caused by overexpression and the GFP fusion). PsvNLS* remains cytoplasmic even in the overexpression and GFP-fusion contexts. (D-F) Expression of wild-type Psv and mutant isoform Psv-NLS* via *C96-Gal4* in developing wings. (D-D’) *C96>GFP* (control) with normal wings (D) and Sens expression (D’). (E-E’) Overexpression of wild-type Psv (*C96>Psv*) showed no effects in wings (E) or Sens expression (E’). (F-F’) Expression of Psv-NLS* (*C96>Psv-NLS\**) displayed partial loss of margin (F) and reduction in Sens expressing cell numbers (F’). (G-I’) Psv and Psv-NLS* interact with ý-catenin/Arm in wing margin specification. Knockdown of *arm* (*C96>arm^IR^*): note partial loss of wing margin (G) and reduction in Sens expressing cells (G’). (H-H’) *C96>Arm^IR^ >Psv*: Co-expression of wild-type Psv rescued the *C96>Arm^IR^* wing defects (H) and restores Sens expression (H’). (I-I’) Co-expression of mutant Psv-NLS* in the *C96>arm^IR^*background (*C96>arm^IR^ >Psv-NLS\**) caused an enhancement of margin defects and also wing tissue loss (I) and markedly increased the loss of Sens expressing cells (I’). Scale bars represent 30μm in salivary glands (B-C), 500μm in wings (G-I), and 50 μm in wing discs and liver cancer cell lines (G’-I’, and L). (J-M) In human cancer cells, Gid8-NLS* blocks Wnt signaling and reduces proliferation (also Suppl Fig. S5). (J) Endogenous Wnt3A expression in cancer cell lines (HCC1395, with HCT116, DLD1 serving as controls). Note high level of endogenous (autocrine) Wnt3A in HCC1395 breast cancer cells. HCT116 and DLD1 cell lines carry mutations in Wnt pathway components, rendering them constitutively active. (K) Relative luciferase assay for Wnt signaling activity in HCC1395 cells, transfected with either LacZ, β-catenin or Gid8-NLS*. Activity level indicates the ratio of TOP-Flash Wnt-reporter firefly luciferase and renilla luciferase activities. Overexpression of β-catenin increases luciferase activity, whereas transfection with the Gid8-NLS* largely eliminates Wnt-signaling activity in HCC1395 cells (\*\**p*<0.01, three independent assays, student’s *t*-test). (L-M) Proliferation assay in HCC1395 cells: Gid8-NLS* transfection suppresses growth. Cells were transfected with LacZ (control) or Gid8-NLS* (L). Note marked reduction in proliferation in Gid8-NLS* transfected HCC1395 cancer cells. Quantified in (M) with \*\**p*<0.01 and \**p*<0.05 (three independent assays, student’s *t*-test).

Next, we asked whether the NLS is required for Psv function in Wg/Wnt-signaling, by expressing the mutant protein, Psv-NLS*, under the control of the *C96-Gal4* driver in developing wing imaginal discs (Fig. 5D-I’). Strikingly, expression of Psv-NLS* in developing wings leads to a loss of wing margin cells and marked reduction in cells expressing the Wg-signaling target Sens (Fig. 5F-F’; note that the equivalent expression of wild-type Psv has no effect on wing development, Fig 5E-E’; compare to wild-type control, Fig. 5D-D’). To confirm its functional interaction in the β-catenin/Arm nuclear translocation context, we compared the effect of wild-type Psv and Psv-NLS* on a mild *arm^RNAi^* knockdown background (see control in Fig. 5G-G’ for *arm^RNAi^*alone). While overexpression of wild-type Psv largely rescued the *C96>arm^RNAi^*defects, both as seen in adult wings and by the expression of the Wg-signaling target Sens (Fig. 5H-H’), co-expression of the Psv-NLS* mutant markedly enhanced wing margin and tissue loss in adults and loss of Sens expressing cells in discs (Fig. 5I-I’). These data indicate that the NLS of Psv is critical for its function in the β-catenin/Arm context.

To address whether this role of Psv is conserved in a mammalian setting, we used Wnt-signaling dependent human cancer cell lines and assayed here the NLS requirement of the human orthologue of Psv, Gid8 (see Fig. S1A for sequence alignment). We tested cancer cell lines that are either addicted to autocrine Wnt-signaling (HCC1395; autocrine Wnt3a expression in HCC1395 cells is shown in Fig. 5J) or those that show aberrant, constitutive Wnt-signaling activity independent of ligand (HCT116 or DLD1; refs^40–43^). Expression of a mutant NLS isoform, Gid8-NLS*, in any of these cell lines, either eliminated (HCC1395, Fig. 5K) or markedly reduced expression of the TOP/FOP Wnt-reporter (Suppl. Fig. S5C-D). Consistently, transfection of the Gid8-NLS* mutant blocked proliferation in these cancer cell lines in a functional growth assay (Fig. 5L-M for HCC1395 cells, and Suppl. Fig. 5C-D for DLD1 and HCT116 cell lines).

Taken together with the above *Drosophila in vivo* data, these results confirm a conserved requirement for Psv/Gid8 in Wnt-signaling in general. Furthermore, these experiments establish that the conserved NLS is functional and critically required for the Wg/Wnt-signaling function mediated by β-catenin/Arm, both in *Drosophila* and human contexts. They further provide evidence that Psv/Gid8-NLS* mutants can serve as inhibitors of Wnt-signaling *in vivo* and mutant cancer contexts.

### Psv affects nuclear localization and signaling activity of stable β-catenin/Arm mutations

As the original premise of this study originated from the observation that the very stable deletion mutant β-catenin isoform, ArmS10 (which cannot bind to IFT140, ref^18^, in contrast to Arm*, which is a point mutation in the priming kinase target site ^44^), is still at least partially capable of nuclear translocation, we asked whether interfering with both the IFT-A complex and *psv* would block ArmS10 nuclear entry and action. We first tested this in the wing margin patterning assay (Fig. 6A-D’), where ArmS10 expression induced many ectopic margin bristles as seen in adult wings (Fig. 6C) and super-numerary Sens expressing cells near the prospective wing margin in wing discs (Fig. 6C’). Strikingly, an RNAi-mediated double knockdown of both *psv* and *ift140* in an ArmS10 background (*C96-Gal4>ArmS10*, *>psv^RNAi^*, > *ift140*^RNAi^) fully rescued the ArmS10 gain-of-function defects (Fig. 6D-D’; similarly, *C96>psv^RNAi^*, >*ift140*^RNAi^ fully suppressed the Arm* phenotype, Suppl. Fig. S6A-C’). This is in contrast to single knockdowns of IFT-A components or Kinesin2 that do not block ArmS10 function noticeably^18^ (note that a *psv* single knockdown suppresses ArmS10 markedly but not fully; Suppl. Fig. S6G-G’). Of note, ArmS10 activity is not fully suppressed by *C96>psv^RNAi^*,> *ift140*^RNAi^ double knockdown, as the wing margin resembles wild-type control and does not show wing margin loss (compare Figs. 6A-A’ and 6B-B’ to 6D-D’), suggesting that some ArmS10 nuclear presence remains.

**Figure 6.**
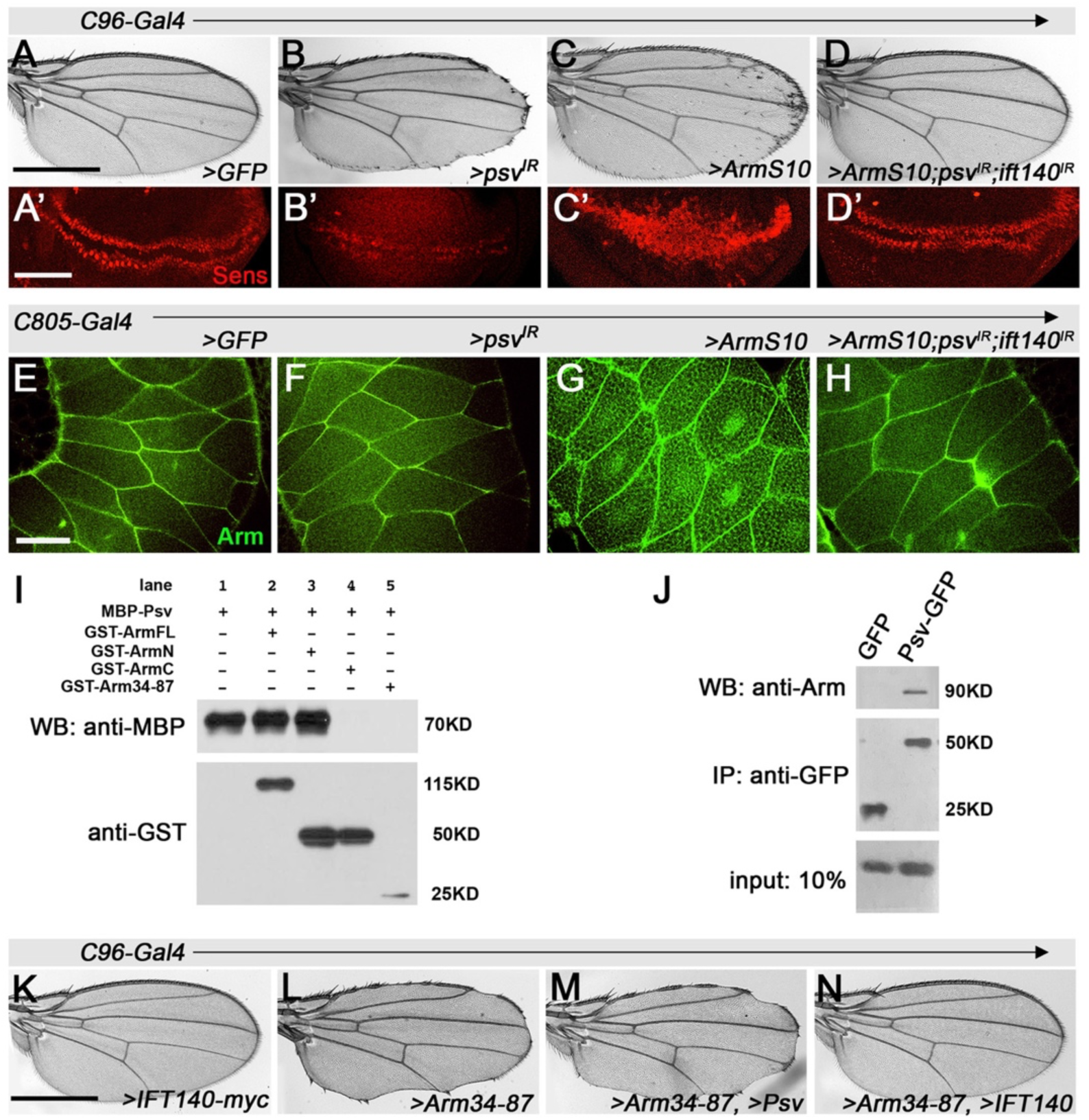
Psv affects stable ArmS10 β-catenin mutation. (A-D’) Wing margin cells induction assay: (A) Control *UAS-GFP* (*C96>GFP*) wing and disc with wild-type Sens expression (red, A′) near D/V boundary. (B-B’) *C96>psv^IR^* (B,B’): note partial loss of margin (B) and reduction in Sens expressing cells (B′). (C-C′) *UAS-ArmS10* (*C96>ArmS10*): note ectopic margin bristle phenotype in wings (C) and supernumerary Sens positive cells (C′). (D-D’) Double knockdown of *psv^IR^* and *ift140^IR^* in *C96>ArmS10* background rescued the defects caused by the stable “constitutively active” ArmS10 mutation. (E-H) Nuclear translocation assay in salivary gland cells (SGs): Without Wg expression (and no LeptomycinB treatment), SGs were stained with anti-Arm, recognizing both endogenous Arm as well as ArmS10 (green). (E) Wild-type control: Arm is in adherens junctions at membranes. (F) No nuclear localization of Arm is detected in SGs of *psv* knockdowns (*C805>psv^IR^*), serving as control. (G) *C805>ArmS10*: expression of ArmS10 reveals that this stable mutant isoform displays nuclear localization (uneven Arm/β-catenin staining caused by mosaic expression of *C805-Gal4*)^54^. (H) *C96>ArmS10, >psv^IR^, >ift140^IR^*: note that double knockdown of both *psv* and *IFT140* largely eliminates nuclear localization of ArmS10, compare to wt control (E). Scale bars represent 700μm for adult wings (A-D, K-N), 50μm in imaginal discs (A’-D’), and 30μm in salivary glands (E-H). (I-J) Molecular dissection of the Psv interaction with Arm/ý-catenin. (I) Direct Psv binding to Arm. MBP-Psv fusion (lane 1 control: input 10%) pulled down by GST-Arm full-length (ArmFL) (lane 2) and GST-Arm N-terminal region (ArmN) (lane 3), but not by GST-Arm C-terminal region (ArmC) or GST-Arm^34–87^ (lanes 4 and 5, compare to lane 1 control). (J) *In vivo* co-IP assay of Psv and Arm from *nub>GFP* (negative control) and *nub>Psv-GFP* wing discs. Protein extracts were co-IPed with anti-Arm. Co-IP and input (10% of lysates used in co-IP assays) were analyzed by blotting with anti-GFP to detect Psv interaction with Arm. (K-N) Functional interactions between Arm^34–87^ peptide and IFT140 or Psv in wing patterning. (K) *UAS-IFT140* (*C96>IFT140*) shows no effect in wild-type, control. (L) *C96>Arm*^34–87^: note partial loss of wing margin cells. (M) Co-overexpression, *UAS-Psv* with *UAS-Arm*^34–87^ does not rescue wing margin loss phenotype caused by *C96>Arm*^34–87^. In contrast, (N) co-expression of IFT140 rescues the dominant negative margin loss phenotype of Arm^34–87^. Note that overexpression of Psv (see Fig. 5E) or IFT140 (K) alone did not affect development.s

We next directly analyzed nuclear translocation of ArmS10 in the salivary gland (SG) assay. The nuclear translocation SG assay is the most reliable measure for nuclear localization. Without Wg expression β-cat/Arm is only detected at adherens junctions (AJs), while when Wg expression is induced nuclear ý-cat/Arm is detected (Fig 1D, see also ^18, 21^). Importantly, the stable ArmS10 mutant isoform is detected in nuclei in the absence of Wg-signaling ^18^. We thus tested ArmS10 behavior in a double knockdown with *psv^RNAi^* and *ift140^RNAi^* in the SG assay (Fig. 6E-H). Strikingly, while single knockdowns did not visibly affect nuclear localization of ArmS10 (ref^18^), the *psv*, *ift140* double knockdown blocked nuclear ArmS10 presence (Fig. 6H; comp. to ArmS10 alone, Fig. 6G).

### Psv and IFT140 physically associate with distinct regions of β-catenin/Arm

To get additional mechanistic insight, we further analyzed the Psv association with β-cat/Arm, and requirement of specific regions in β-cat/Arm mediating this physical interaction (Fig. 6I). Direct *in vitro* pull-down assays with Psv and β-cat/Arm established that full-length β-cat/Arm protein as well as an N-terminal β-cat/Arm fragment bind to Psv (Fig. 6I). This was also confirmed *in vivo* with co-immunoprecipitation of Psv-GFP and endogenous Arm protein (Fig. 6J). Importantly, direct interaction assays (Fig. 6I) established that the Psv interaction with β-cat/Arm was not mediated by the N-terminal Arm region, Arm^34–87^, previously shown to be necessary and sufficient for interaction with IFT140 ^21^. Taken together these data suggest that β-cat/Arm interacts with Psv and IFT140 through neighboring (non-overlapping) sequences in its N-terminal region.

To get further functional insight, we tested the relationship of Arm^34–87^, which acts as a dominant negative for Wg/Wnt-signaling by interfering with IFT140 binding to endogenous β-cat/Arm ^21^. According to the dominant negative behavior of Arm^34–87^, overexpression of IFT140 in the Arm^34–87^ background rescued its effect to wild-type (Fig. 6K, L and N). We thus asked whether Psv, which is similarly critical for nuclear β-cat/Arm translocation, can also functionally interact with the Arm^34–87^ dominant negative effect. Strikingly, and in contrast to IFT140, overexpression of wild-type Psv (which itself showed no defects in a wild-type background, Fig. 5E) did not rescue the phenotype caused by Arm^34–87^ expression (Fig. 6M, compare to 6L, showing the Arm^34–87^ expression defects).

Taken together, all these data are consistent with the model that IFT140 and Psv are both required for nuclear β-cat/Arm translocation, and that they act in parallel, not redundantly or interchangeably (see Discussion).

## Discussion

In this study, we define the function of a novel, conserved protein, Pasovec (Psv, Gid8 in humans), within the canonical Wg/Wnt-pathway and its associated requirement for nuclear translocation of β-catenin/Arm. Our data reveal that Psv (and its human orthologue Twa1/Gid8, suggested to interact with RanBPM ^23^), which contain a nuclear localization sequence/NLS can directly bind to both the N-terminal region of β-catenin/Arm and to IFT140. β-catenin is a multifunctional protein, which (i) associates with AJs and links these to the actin cytoskeleton, and (ii) is required in canonical Wg/Wnt-signaling, where it acts as the key nuclear effector. Although entry of cytoplasmic β-catenin/Arm into the nucleus is critical in Wnt-signal transduction, β-catenin/Arm does not contain an NLS. We show that the NLS of Psv is required for β-catenin/Arm nuclear localization and, importantly, *Drosophila* Psv or the human orthologue harboring a mutant NLS serve as dominant inhibitors of Wnt-signaling. As Psv is also found associated with IFT140 within the IFT-A complex (which is itself required for nuclear translocation of β-catenin/Arm ^18^) in the cytoplasm independent of Wnt-pathway activation, it appears to be a critical component of the machinery promoting nuclear β-catenin/Arm translocation. Taken together, our findings indicate that Psv and IFT-A work together to promote nuclear β-catenin/Arm localization upon Wg/Wnt-signaling activation.

### Psv/Gid8 are required for Wg/Wnt signaling

The failure to identify Psv earlier as a component of Wg/Wnt-signaling might be surprising. It is encoded by a small gene and thus could have been missed in the genetic screens for patterning mutations, which were not fully saturating^45^. Many of the subsequent identifications of critical components of the canonical Wg/Wnt-pathway were associated either with genetic screens using modifier assays in *Drosophila* or by molecular screens and biochemical assays in *Xenopus* (rev. in ^46–48^). For example, in *Drosophila,* a gain-of-function (GOF) mis-expression phenotype of Wg in the eye was screened for suppressors and this approach identified the TCF-family transcription factor, *pangolin*, and associated transcriptional co-factors Legless and Pygopus (refs^49, 50^). Similarly, GOF molecular screens identified several Wnt-pathway components in the frog system^46–48^. While the function of Psv/Gid8 would have likely been missed in the GOF assays in *Xenopus*, it could have been found in the suppressor screen in the *Drosophila* eye^49, 50^. Nonetheless, no screening approach – genetic or molecular - is likely to identify all factors and components of a process, and as such it is expected that new pathway components, in any context, can still be discovered. As such genetic approaches in model organisms, like *Drosophila*, remain a very valuable tool to complement the vast data coming from molecular ‘omics’-approaches.

### What is the functional relationship of IFT-A and Psv in Wnt-signaling?

The definition and mechanistic dissection of the cytoplasmic role of IFT-A in Wnt-signaling and nuclear β-catenin/Arm localization has been a breakthrough to start understanding the process of nuclear β-catenin/Arm translocation^18^. However, the observation that the stable mutant ArmS10 isoform, which is lacking the N-terminal region required for IFT140/IFT-A binding, can still enter the nucleus suggested that other factors must be participating in β-catenin/Arm nuclear import ^18^. Strikingly, we demonstrate here that in a double mutant scenario, with both IFT-A and Psv impaired, nuclear localization of ArmS10 is indeed inhibited. At first, this might suggest that the IFT-A complex and Psv share partially redundant roles in the process. However, in mutant clones of either *psv* (this study) or *IFT140* (ref^18^), Wg-signaling targets, like Sens, are not expressed, indicating a non-redundant function for both factors. Similarly, *psv* mutant embryos strongly resemble the phenotypic features of *arm* mutant embryos that affect Wg-signaling, and in the salivary gland nuclear localization assay knockdown of both block nuclear β-catenin/Arm translocation. Moreover, (over)expression of the N-terminal region of β-catenin/Arm, Arm^34–87^ or human β-catenin^24–79^, which is necessary and sufficient to bind to IFT140, inhibits very effectively Wnt-signaling during *Drosophila* development and in human cancer cell lines by blocking nuclear β-catenin/Arm translocation, including that of stable β-catenin/Arm mutations that still bind to IFT-A (ref^21^). Taken together, these results suggest a non-redundant function. The observation that Psv also binds to IFT140 and β-catenin/Arm is consistent with the notion that Psv acts together with the IFT-A complex. Importantly, Psv binds the β-catenin/Arm N-terminal region in sequences adjacent to and non-overlapping with the Arm^34–87^ fragment, which binds IFT140.

The outlier in the logic of non-redundant functions of IFT140/IFT-A and Psv is again the ArmS10 deletion. However, ArmS10 might behave somewhat differently as it would escape cytoplasmic retention mediated by N-terminal sequences due to the nature of its deletion. In this context, it is noteworthy that, while the effects of ArmS10 are strongly suppressed by the double knockdown of *IFT140* and *psv*, the suppression *in vivo* does not drop below “wild-type levels”, as represented by a normal wing margin (see Fig. 6D), suggesting that some ArmS10 still escapes the suppression, providing sufficient signaling levels to maintain close to wild-type levels. As ArmS10 is extremely stable, it is possible that low background levels of translocation, not employing the regular mechanism, are sufficient to maintain some signaling.

### Psv appears to provide the critical NLS within the IFT-A/Psv/β-catenin/Arm complex

While IFT-A is critical for the nuclear translocation of β-catenin/Arm and its association with the cytoplasmic MT network, it lacks an NLS. Thus, within the IFT-A/Psv/β-catenin complex described here, Psv likely serves as the key factor for providing the function of an NLS. Accordingly, mutating the NLS of Psv blocks the nuclear translocation of β-catenin/Arm. The observation that overexpression of this mutant form can interfere with the process, in analogy to a dominant negative interference tool, provides additional evidence for its critical requirement besides the IFT-A/Kinesin2 complex. In summary, nuclear translocation of β-catenin/Arm can be effectively interfered with from both sides, either (over)expressing (1) the N-terminal peptide of β-cat/Arm, Arm^34–87^ or human β-catenin^24–79^, which blocks its interaction with IFT-A; or (2) a Psv-NLS mutant, which interferes via its non-functional NLS. While the NLS “requirement” of Psv resembles general (canonical) nuclear import mechanisms, the required IFT-A involvement maintains the unique mechanism required for β-catenin/Arm nuclear import.

## ACKNOWLDGEMENTS

We THANK Jun Wu, Ursula Weber and all other Mlodzik lab members for helpful input and discussion. We are grateful to H. Bellen for the Sens antibody, K. Cadigan for the Arm* fly strains, T. Avidor-Reiss for IFT-A strains, and KW. Choi for the Kinesin construct. We also thank the *Drosophila* stock centers (Bloomington Indiana, NIG, and VRDC) for many fly strains, the Developmental Studies Hybridoma Bank for antibodies, and FlyBase for information about *Drosophila* stocks and reagents. We are most grateful to Stuart Aaronson for kindly providing all human cell lines used in this study. We thank Robert Krauss, Prashanth Rangan, and Ursula Weber for very helpful comments on the manuscript. We would like to thank the ISMMS Microscopy CoRE, where confocal microscopy was performed, which was in part supported by the Tisch Cancer Institute P30 CA196521 grant from the NCI. This work was supported by NIH/NIGMS grant R35 GM127103 to MM.

## AUTHOR CONTRIBUTIONS

LTV and MM designed the study, LTV performed all experiments and developed experimental tools. MM and LTV analyzed the data and wrote the manuscript. MM provided funding for the study.

## MATERIALS AND METHODS

### *Drosophila* stocks and Flybase information access

The Gal4/UAS system was used for expression of RNAi constructs (sometimes in combination with *UAS-Dcr2*) and other transgenes. Gal4-driver for wing margin during wing development was *C96-Gal4* expressed around the dorsal-ventral compartment boundary (Suppl Fig. S1), and for salivary glands *C805-Gal4* was used. All crosses were set up at 25°C (or at 29°C where indicated). We frequently access information in FlyBase ^51, 52^ to get details on many of the *Drosophila* stocks used throughout this study, these are all listed in the Key Resources Table below.

### Generation of *psv* mutants

The primers (listed under Key Resources) were inserted as gRNA into pBFv-U6.2B. This construct was then injected in the TBX-0002 *y^1^v^1^ P{nos-phiC31\int.NLS}X; attP40* Drosophila stock and crossed with CAS-002 *y^2^ cho^2^ v^1^ P{nos-Cas9, y+, v+}1A/FM7c, KrGal4 UAS-GFP* for generation of CRISPR mutant alleles. Mutants were selected by non-lethal genotyping and amplification of the targeted region by PCR. To make the *psv* C-term deletion, a gRNA targeting sequence (key resource table) was cloned into pCFD3 and the *psv*-targeting gRNA transgenic fly was made by microinjection. The *psv*-gRNA strain was then crossed with *nos*-Cas9 flies, the F1 progeny from this cross were microinjected with a HDR donor, and the F2s were screened for the *psv* C-term deletion. All these steps were performed by BestGene.

The Gid8-NLS*mutant was constructed using the In-Fusion HD Mutagenesis Kit from Clontech (Takara) to generate the respective point mutations. PCR primers are indicated in Resources table.

### Transgene construction

To generate transgenic flies, *psv* was amplified by PCR using DGRC LD23131 cDNA (for *psv*) and cloned into *pUAS-attB* and *pUAS-attB-GFP* vectors (VK1, second chromosome 2R 59D3) using EcoRI and XbaI sites. The injection step and fly screening were done by Bestgene.

### Generation of anti-Psv antibody

The entire protein coding sequence of Psv was amplified and cloned into pGEX-4T1 through *Eco*RI and *XbaI* sites. GST-Psv fusion protein was expressed in the BL21 strain by isopropyl β-D-1-thiogalactopyranoside induction, and purified protein was used to immunize rabbit and rat (MSBI antibody company).

### Cuticle preparation and wing mounting

For cuticle preparation, dechorionated embryos were mounted in Hoyer’s medium^53^, incubated at 60°C overnight and viewed with a Zeiss Axiolmager microscope. Wings from adults were removed, incubated in PBS with Triton detergent, and mounted on a slide in 80% glycerol in PBS, and imaged using Zeiss Axioplan microscope. All adult images were acquired using Zeiss Axiocam color-type 412-312 camera and Zeiss axiocam software.

### Immunostaining and histology

Imaginal discs were dissected at 3^rd^ instar larval stage in PBS and fixed in 4%PFA in PBS. Discs were washed 2 times in PBS, 0.1% Triton-X100 (PBT), incubated in primary antibodies o/n at 4°C. After washing in PBT, incubation with secondary antibodies was at RT for 2hrs. Samples were mounted in Vectashield. Wing disc images were acquired with a confocal microscope (20X-40X, oil immersion, Leica SP8 or Zeiss LSM980 system). Images were processed with ImageJ (National Institutes of Health) and assembled in Photoshop (Adobe).

Salivary glands were dissected at 3^rd^ instar larval stage in PBS and treated with 0.1% Leptomycin B (Sigma) for 5-10 min before fixation in PBS, 4% PFA. All subsequent steps were as described for wing imaginal discs.

### GST pull down

For GST pull-downs, IPTG-inducible E. coli R2 cells (BL21) were transformed with plasmid constructs for MBP fusion proteins MBP-Kap3, MBP-Klp64D, MBP-Klp61F, MBP-IFT140, MBP-Arm, and MBP-Psv, as well as GST-fusion proteins GST-Arm, GST-ArmN, GST-ArmC, GST-Arm^34–87^, and GST-Psv. Fusion proteins were purified from bacterial lysates. An equal amount of blocked glutathione Sepharose 4B beads with GST, GST fusion protein or beads alone were incubated with lysates containing MBP-fusion proteins O/N at 4°C. After several washes with pull-down buffer (20 mM Tris pH 7.5, 150 mM NaCl, 0.5 mM EDTA, 10% glycerol, 0.1% Triton X 100, 1mM DTT, and protease inhibitor cocktail), sample buffer was added, beads were boiled, and protein were resolved by SDS-PAGE. For Western blotting, proteins were transferred onto nitrocellulose, blocked in 5% skim milk and incubated with primary goat anti-GST or rabbit anti-MBP antibody. Protein bands were visualized using Immobilon Forte Western HRP Substrate kit.

### Immunoprecipitation

Lysates from 30 wing imaginal discs of *nub>IFT140myc, nub> IFT144myc; Psv or GFP* were precleared by incubating with protein A-Agarose beads for 1hr at 4°C followed by centrifugation. A-Agarose beads were immune-precipitated with specific antibodies at 4°C for 1hr. Polyclonal anti-GFP antibody or anti-Psv were used. Immunoprecipitates were resuspended in SDS sample buffer, boiled for 5 min, separated by SDS-PAGE, and transferred to nitrocellulose for immunoblotting. Protein was detected by Immobilon Forte Western HRP Substrate kit.

### Cell lines and culture conditions

Wnt3a-expressing L-cells and control L-cells were grown in Dulbecco’s Modified Eagle’s Medium (DMEM) supplemented with 10% fetal bovine serum (FBS), and 1% penicillin/streptomycin. Wnt3A and control L-cells were prepared as described by a protocol provided by ATCC. Human cancer cell lines DLD1, HCT116 were grown in DMEM supplemented with 10%FBS. HCC1395 cell line was grown in RPMI 1640 medium supplemented with 10% FBS. All cells were cultured at 37°C in 5% CO_2_. All cell lines were kindly provided by Stuart Aaronson.

### Luciferase reporter assay for β-catenin activity (TOP/FOP reporter assay)

Human full-length β-catenin, Gid8-NLS* or lacZ were cloned into pcDNA4/TO plasmids. For all cancer cell lines, cells were seeded at a density of 4×10^5^ cells/ 12-well plate one day before transfection. Cells were then transfected with plasmid constructs (LacZ, full-length β-catenin or Gid8-NLS*: 1 *μ*g each) together with 2*μ*g pGL-TOP or its negative control vector pGL-FOP. For HCC1395, HCT116 or DLD1 cell lines (gifts from Stuart Aaronson), after 24h, Wnt3A condition medium was added to the cells and incubated for additional 24h. The treated cells were then washed and lysed using 20 *μl* luciferase lysis buffer per well, and luciferase activities were performed and measured in 96-well plates using Dual-Luciferase Reporter Assay system according to the manufacturer’s protocol. The measurement was conducted on Synergy MX luminometer. Experiments were carried out in triplicate and repeated at least three times, shown as the mean ±SD of the ratio between the TOP/FOP.

### Cell proliferation assay

The cell proliferation assay was determined by 3-(4,5-dimethylthiazol-2-yl)-2,5-diphenyl-tetrazolium bromide (MTT) assay (Sigma). All cancer cell lines were seeded in 24-well plates (2×10^4^ cells/well). After 24hr, cells were transfected with full length β-catenin or Gid8-NLS*. Absorbance at 570nm was read on a microplate reader. All assays were performed in triplicate.

## QUANTIFICATION AND STATISTICAL ANALYSIS

### Quantitative analysis of wing discs

Wing imaginal discs staining were processed using Image J. To establish an appropriate quantification of expression/staining signal, Sens intensity was normalized by subtracting the signal in the Sens negative (-) cells from signal obtained in the Sens positive region, “(+) cells”. Mean measurements were plotted for 10 wing discs. For statistical analyses, a two tailed test was performed on normalized mean intensity measurements to compare genotypes.

## KEY RESOURCES TABLE

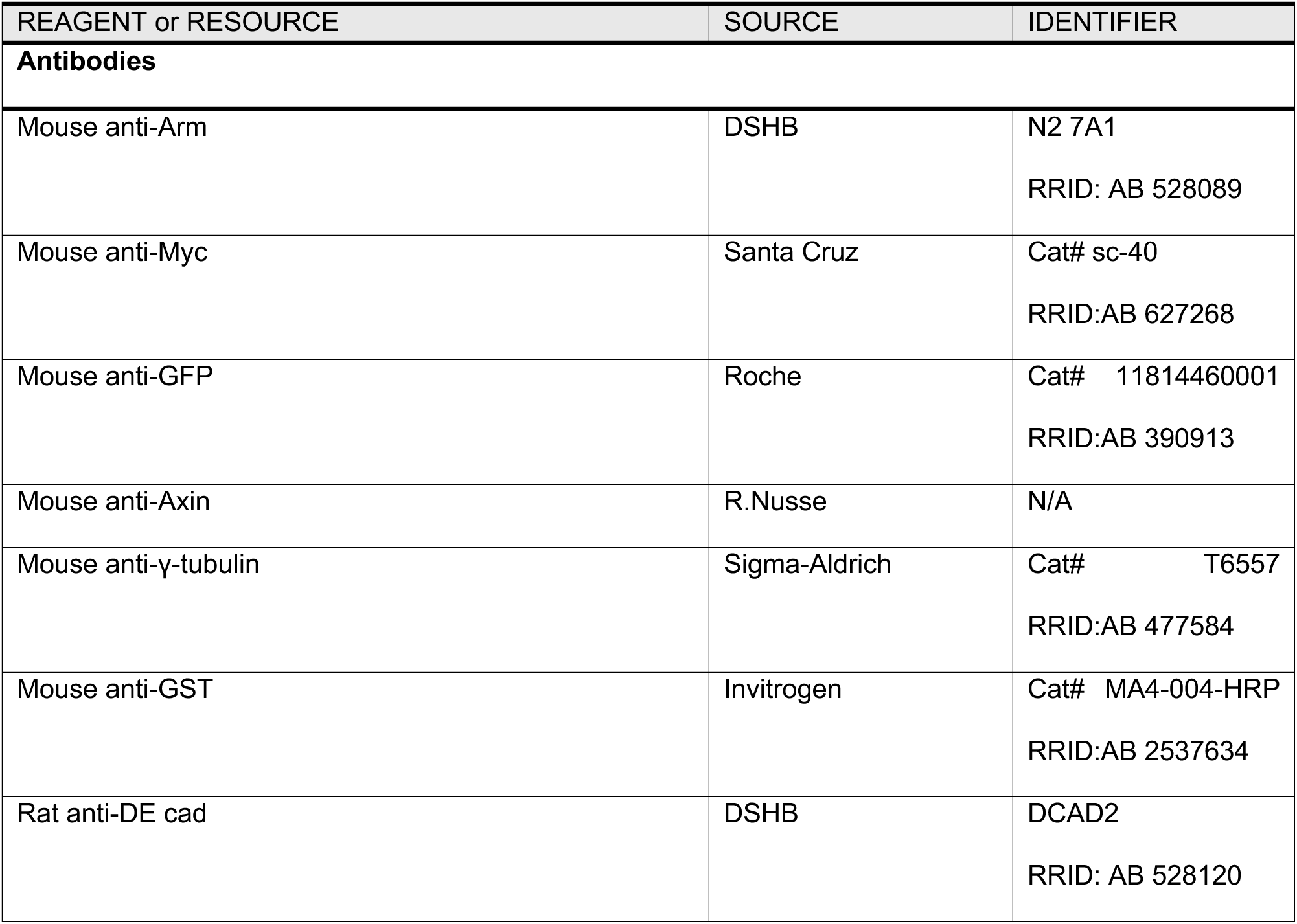
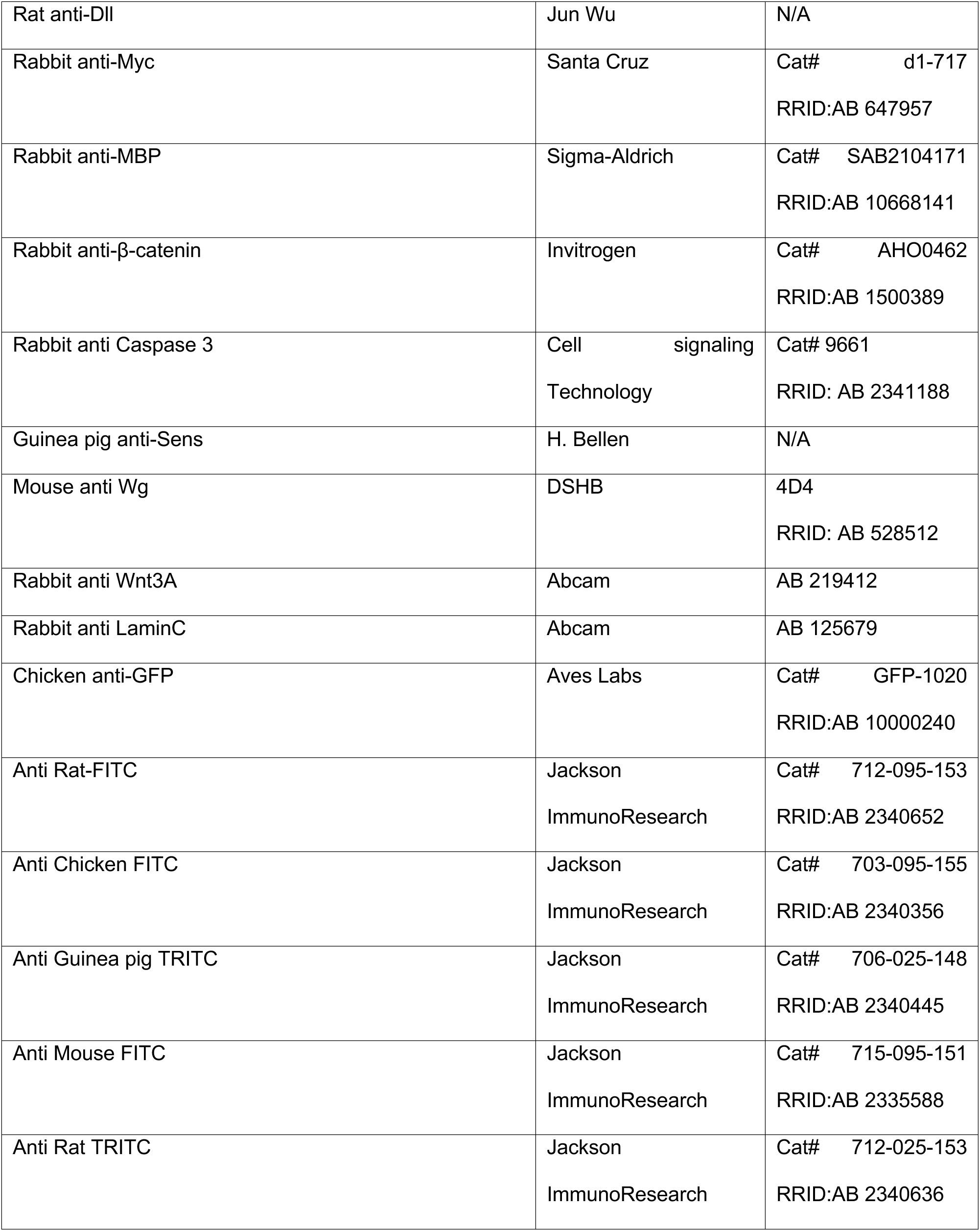
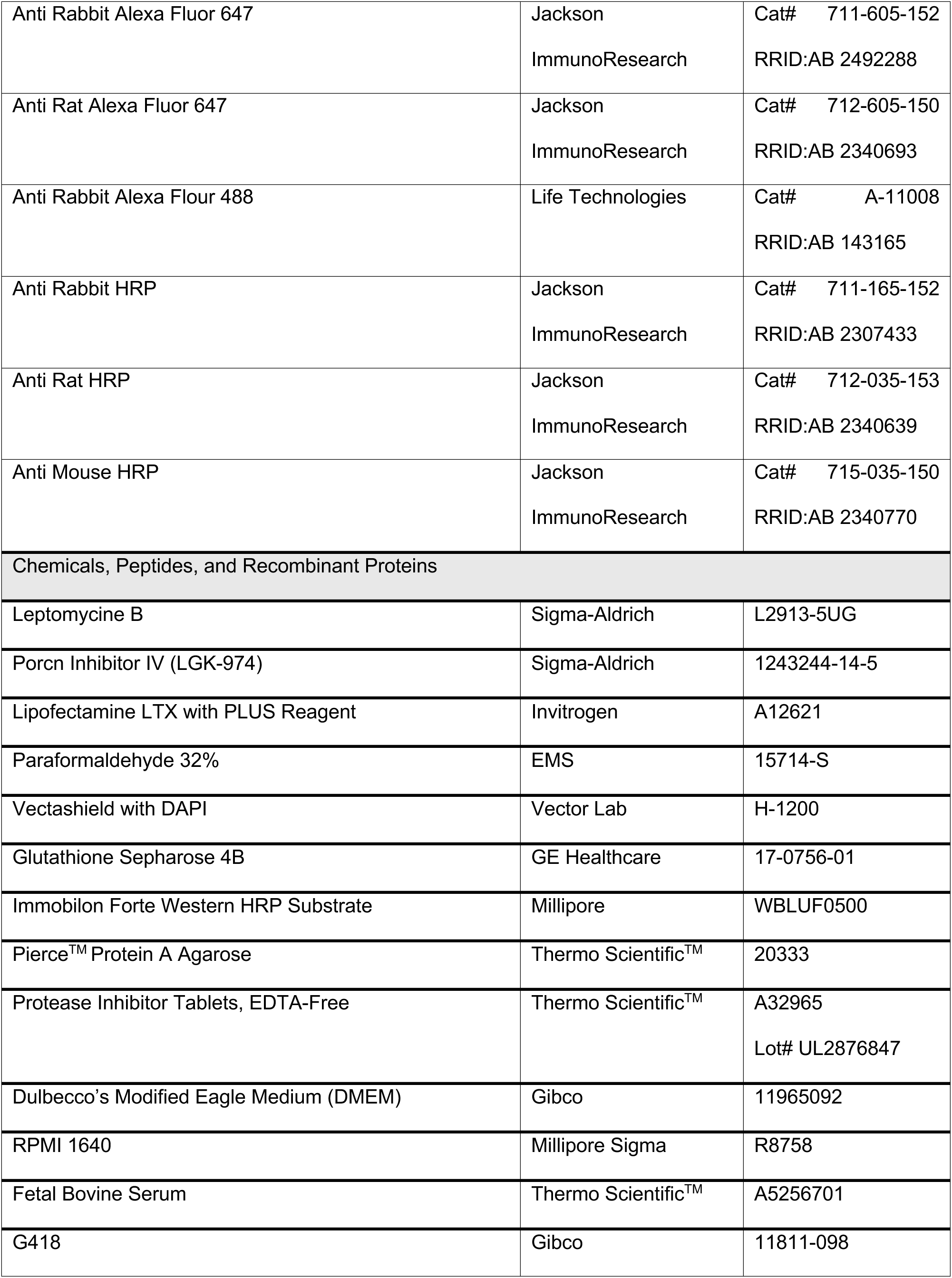
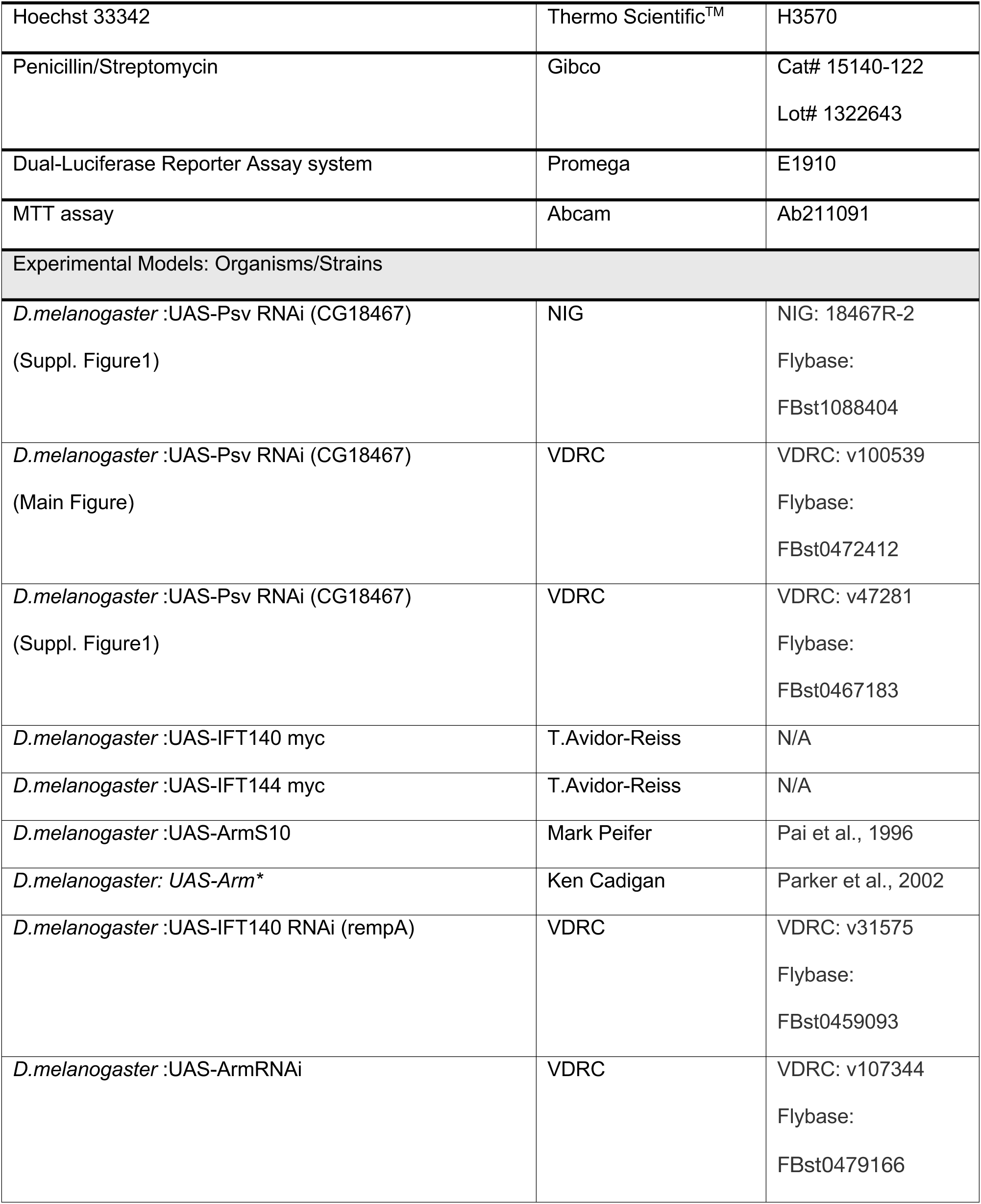
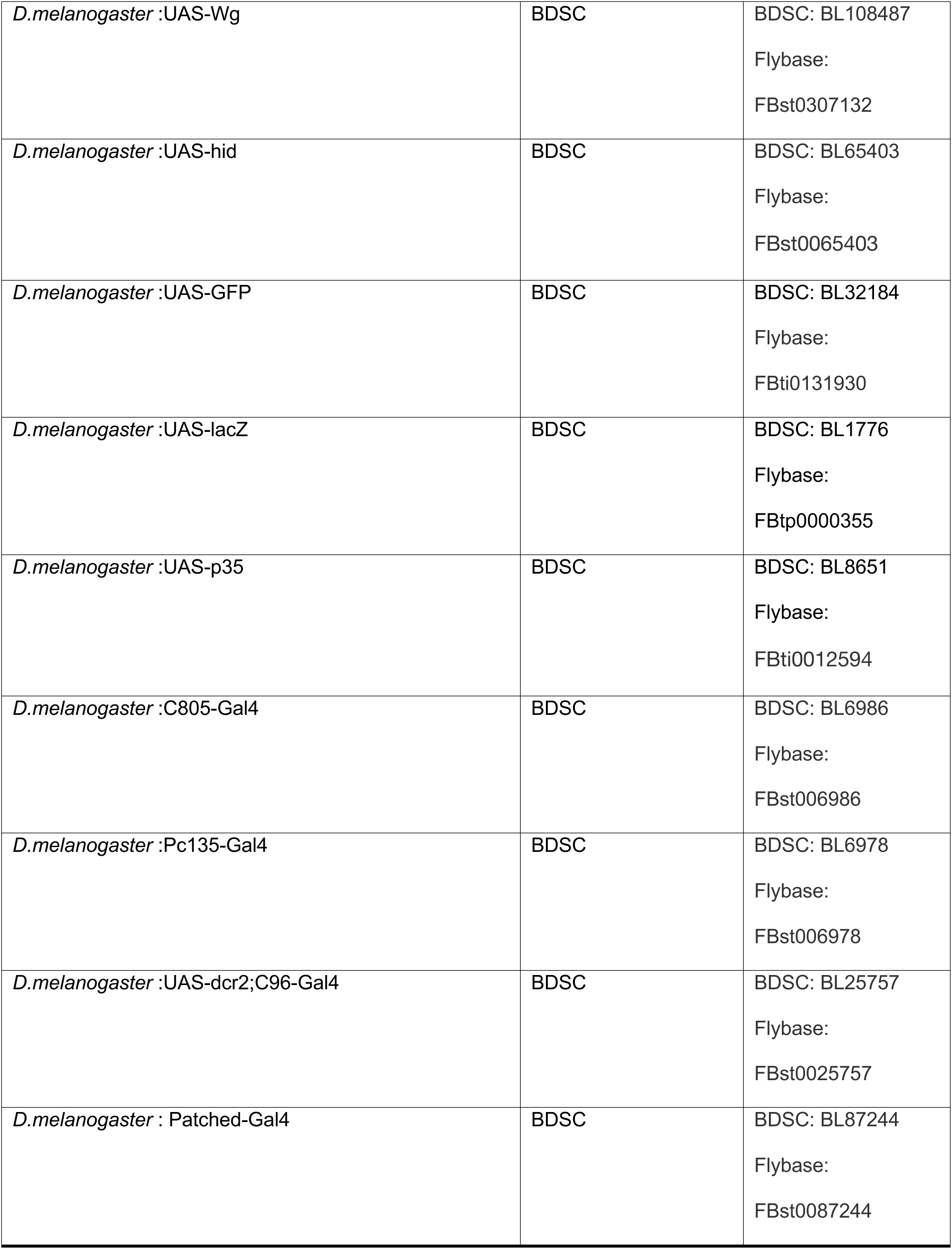
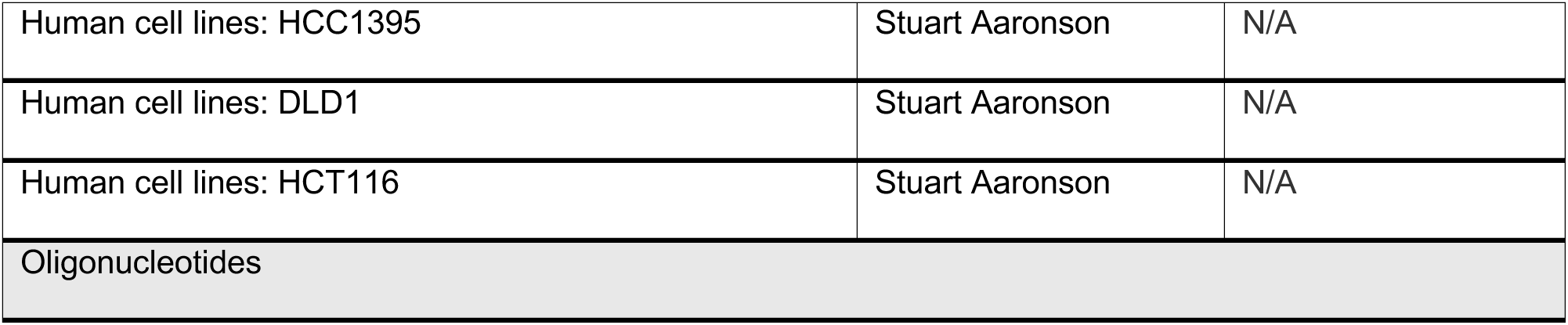
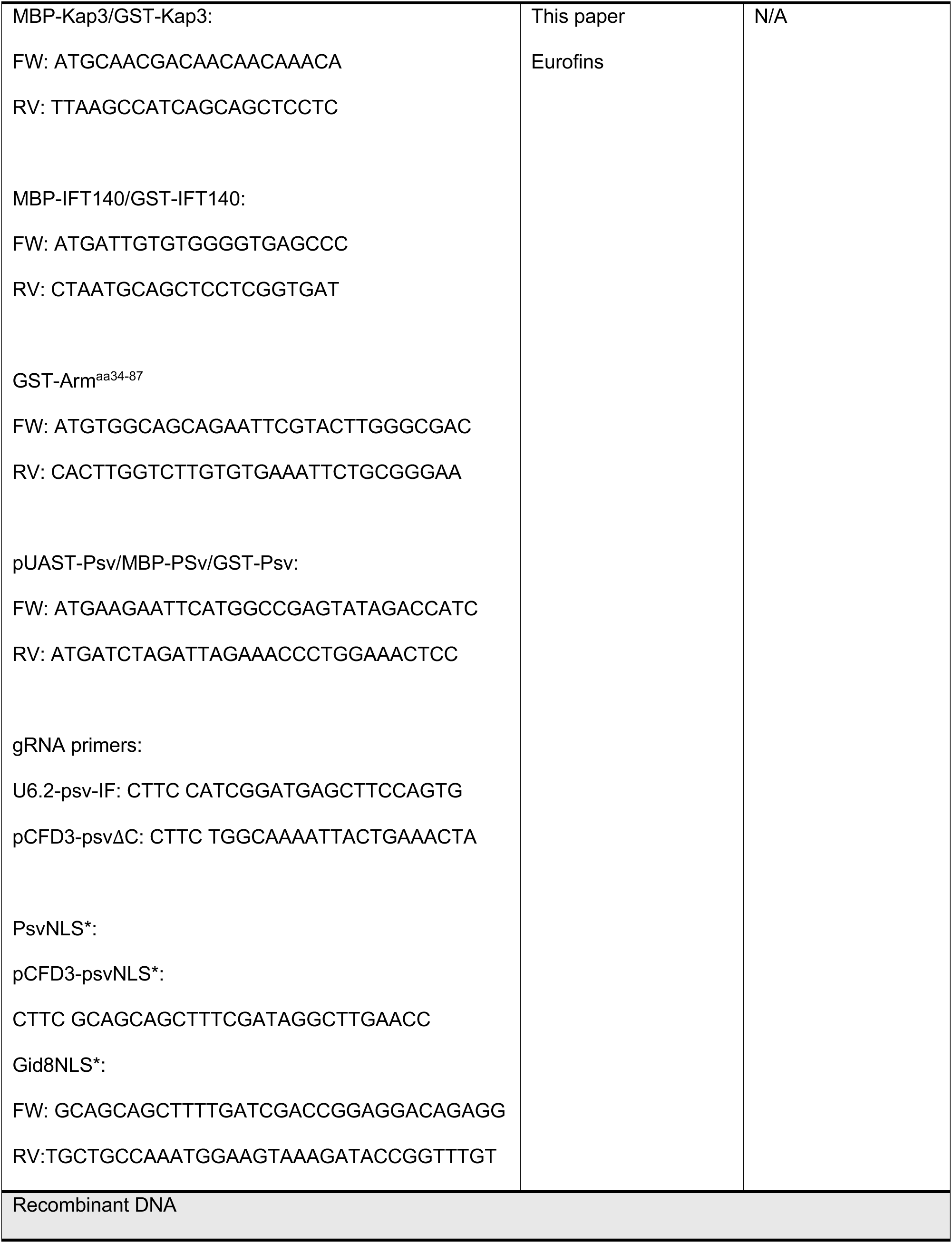
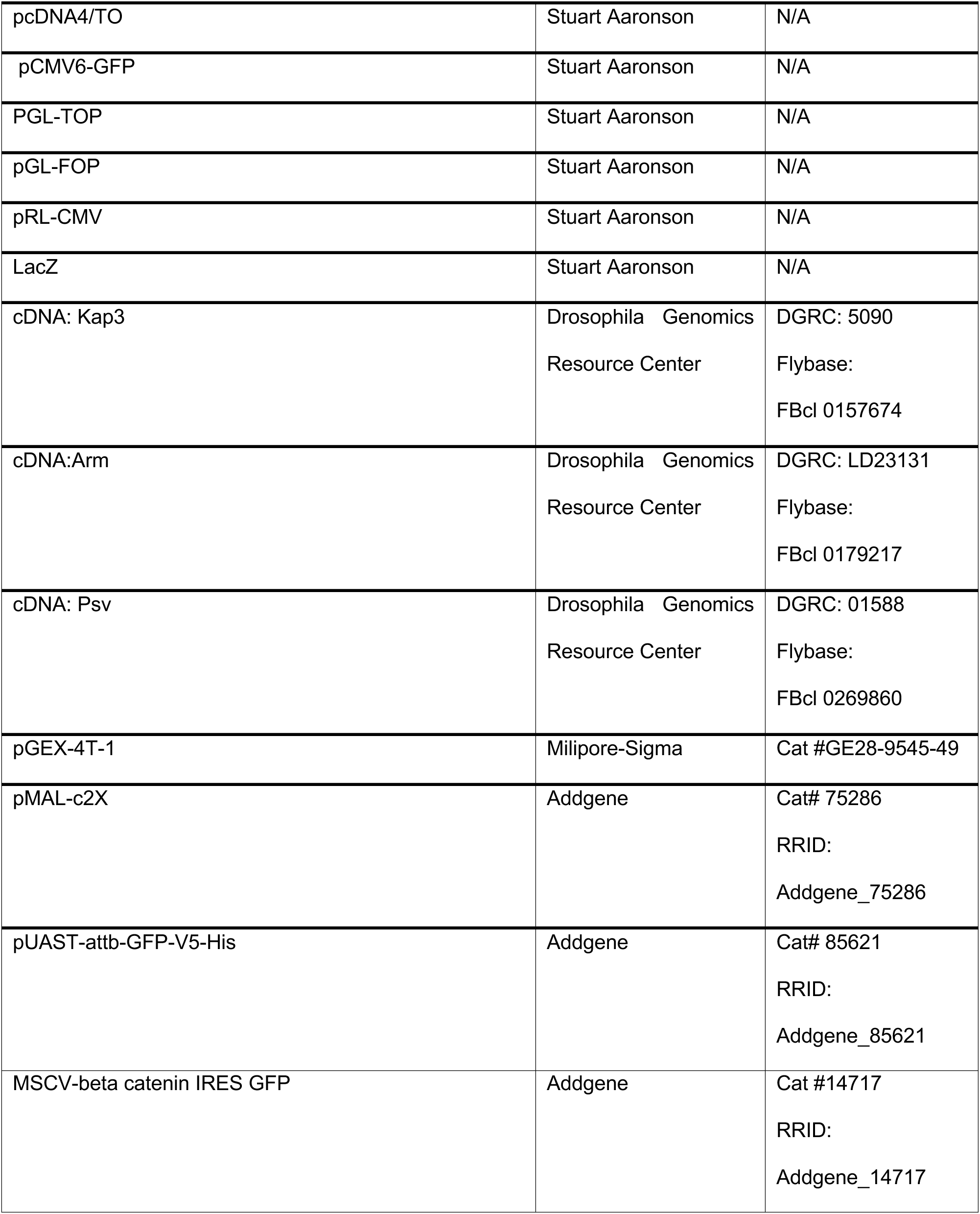
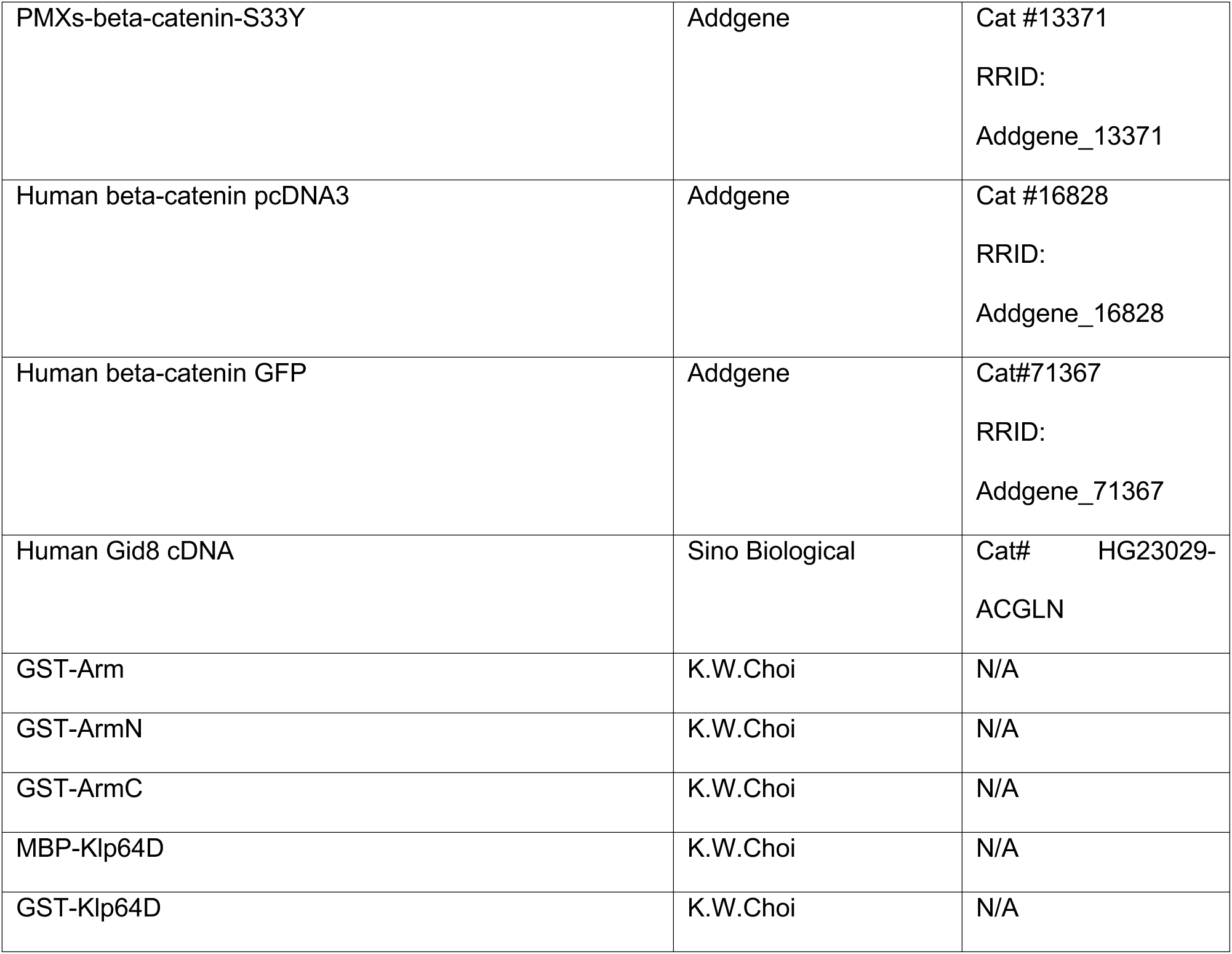

