## Supplemental Data for "A novel conserved protein associates with the IFT-A complex to mediate nuclear translocation of β-catenin in Wg/Wnt-signaling"

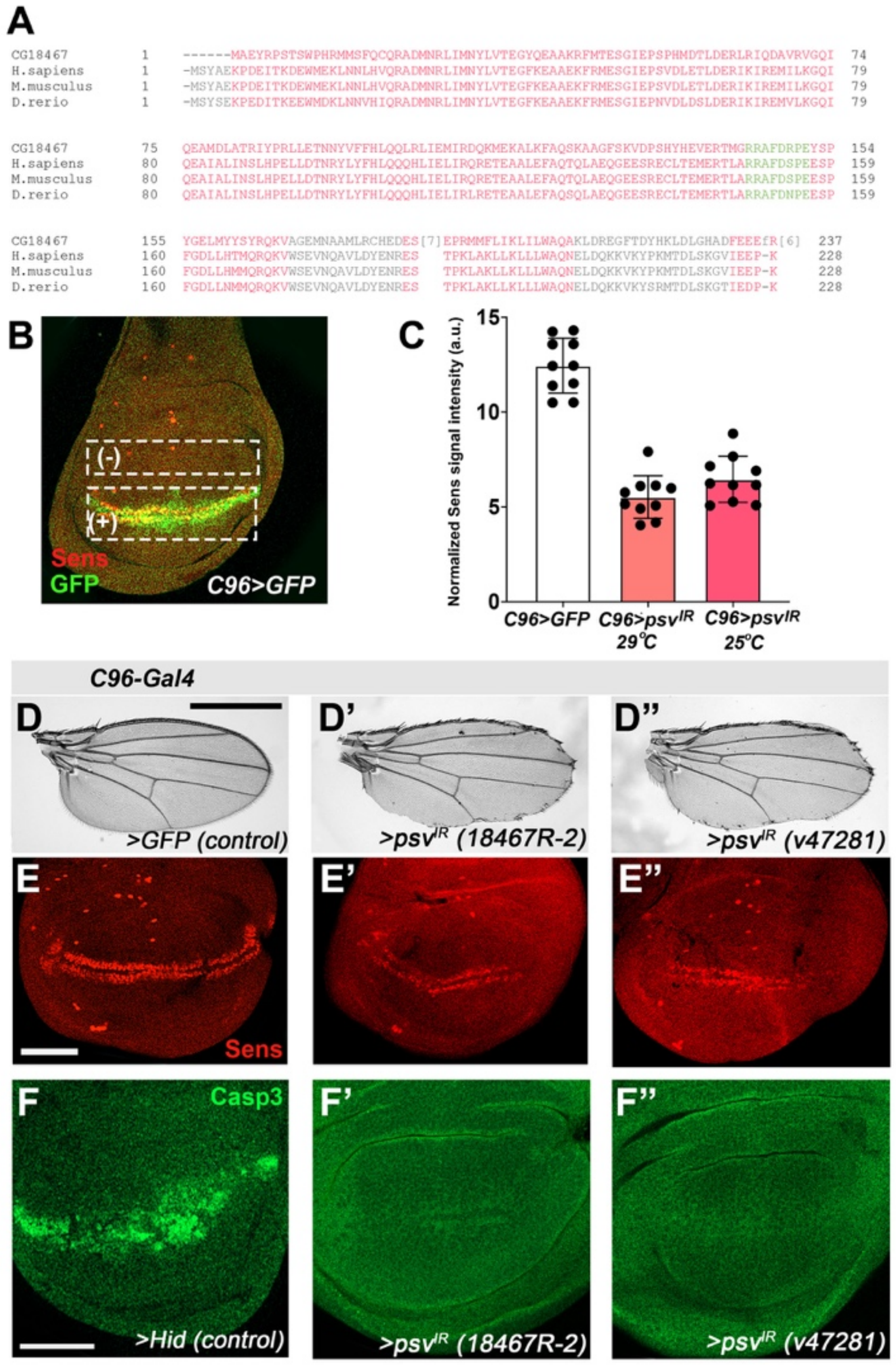


**Figure S1. Psv is required for wing margin development and Wg target gene expression.**
**(A)** Sequence alignment of *Drosophila* CG18467/Psv with its orthologs human, mouse, and zebrafish, generated using NCBI blast. Highly conserved residues are highlighted in red and the nuclear localization sequence (NLS) is shown in green.
**(B)** Expression domain of the *C96-Gal4* driver as shown with UAS-GFP, *C96>GFP*, completely overlapping the Sens expression region (red, GFP in green). Regions of Sens expression quantification are shown in boxed areas (white rectangles indicated by "+" for Sens positive region, and "-" for Sens negative/control region). To establish an appropriate quantification of the Sens signal strength, we

normalized Sens signal intensity in "+" ROI by subtracting the signal in the Sens negative (-) ROI from the signal obtained in Sens "(+) cells".

**(C)** Quantification of Sens staining as affected by *psv* RNAi in Figure 1. *C96>GFP* wing disc serves as the positive (wild-type) control, while *C96>psv RNAi* results in partial loss of Sens expression.

**(D-E")** Effects of two additional independent *psv* RNAi transgenes at 25°C driven by *C96-Gal4* (compare to main Figure 1). *C96>GFP* control show normal adult wing (**D**) and Sens expression in wing discs (**E**), whereas the two *psv* RNAi transgenes result in partial loss of wing margin (**D, D"**) and reduction in Sens expressing cells (**E', E"**).

**(F-F')** *psv* knockdown does not induce cell death as determined with anti-Caspase3 (Casp3) staining (green). (F) shows positive control of Hid expression in the C96-Gal4 domain, note abundant Casp3 staining. Knockdown of *psv*, mediated by two independent *psv*-IR transgenes does not induce Casp3. Scale bar in **D-D"** represents 800μm, 50 μm in **E-E"** and 30 μm in **F-F"**.

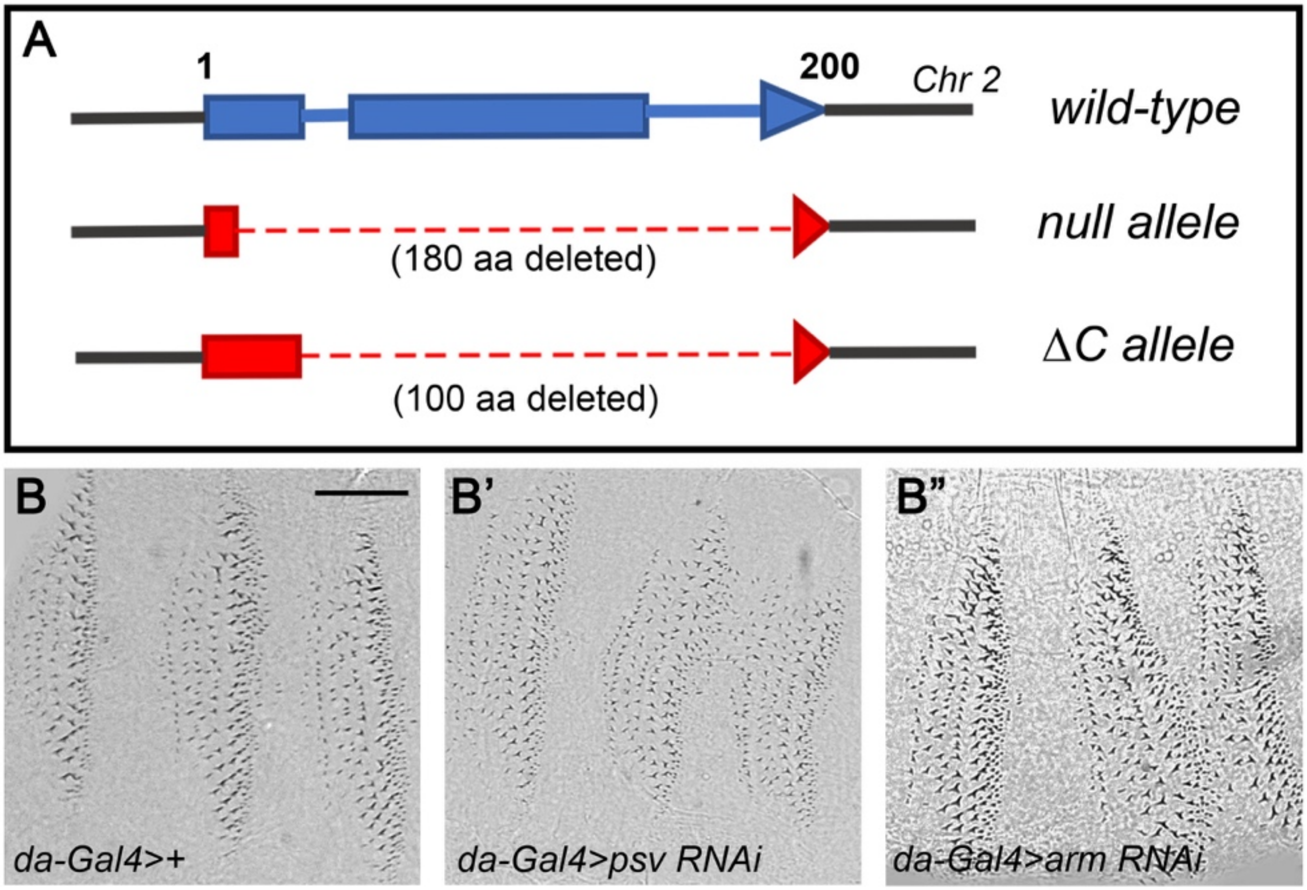


**Figure S2. Knockdown of *psv* affects embryo patterning similar to *arm* knockdown**.
**(A)** Two deletion mutations lacking most of the *psv* coding sequence (null mutation) and the C terminal region (ΔC).
**(B-B'')** Knockdown of psv using *daughterless-Gal4* (*da-Gal4*) results in the formation of ectopic denticles belts at the expense of naked cuticle **(B')**. These defects resemble the phenotype of knocking down *arm* (*da-Gal4>arm RNAi*) (**B''**, compare to wildtype (**B**). Scale bar in **B** indicates 80μm.

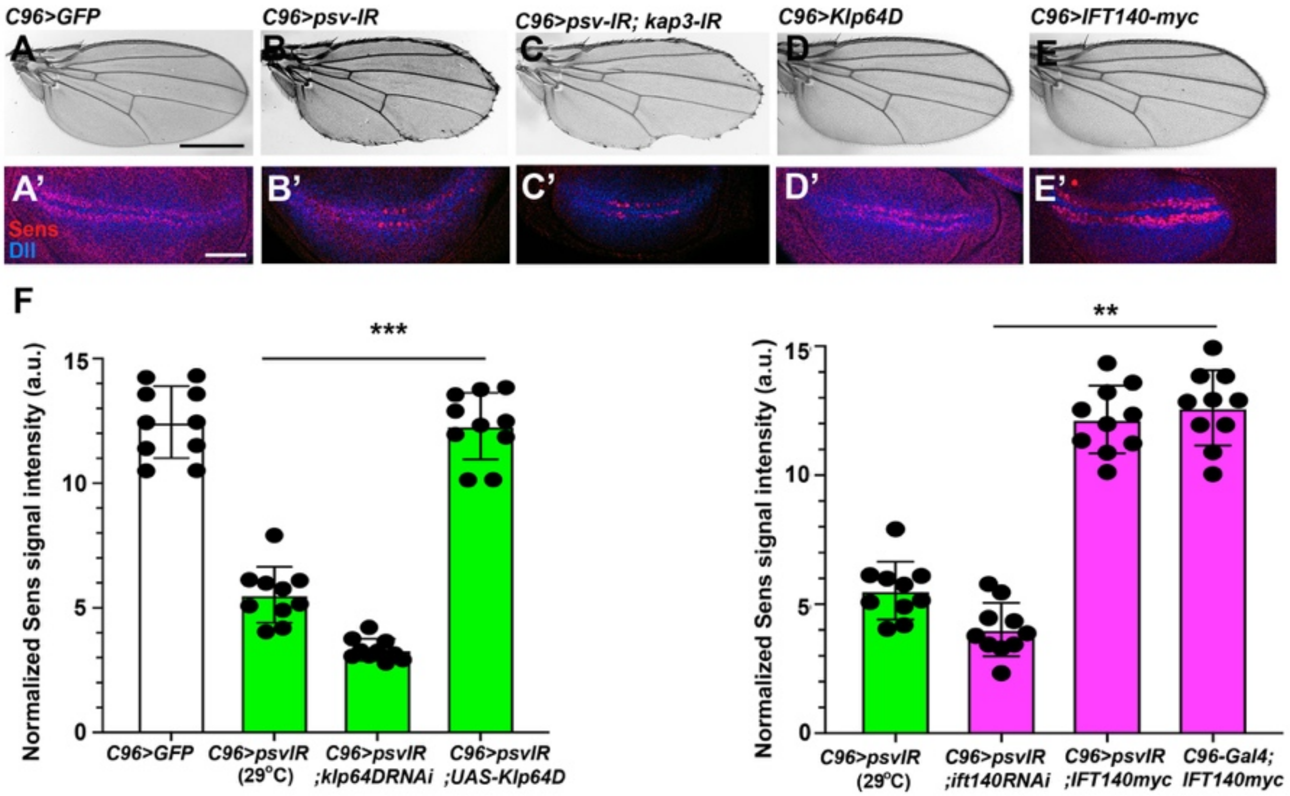


**Figure S3. Interaction between *psv* and *kap3*.**

**(A-C)** Functional interaction between *psv* and *kap3* during wing patterning. *UAS-GFP* (*C96>GFP* control wing (**A**) and equivalent control wing disc with wild-type Sens (red) and Dll levels (blue) expression near DV boundary (**A'**). (**B-B"**) *C96>psv$^{IR}$* wing and wing disc. Note partial loss of margin (**B**) and partial loss of Sens and reduction of Dll (**B'**), consistent with adult wing defects. (**C-C'**) *psv* and *kap3* double knockdown. *C96>psv$^{IR}$, >kap3$^{IR}$*: note increased wing margin loss (**C**), and increased reduction in Sens and Dll expression (**C'**). Scale bars in **A** (for **A-E)** and **A'** (for **A'-E'**) represent 700 μm and 50 μm, respectively.

**(D-E)**. Overexpression of Klp64D/Kinesin2 (**D-D'**) or IFT140-myc (**E-E'**) showed normal wing phenotypes (**D, E**) and normal expression of Sens and Dll (**D'-E'**). All genotypes were reared at 25°C.

**(F)** Quantification of normalized Sens staining level from panels in main Figure 3 as described in Suppl. Fig S1.

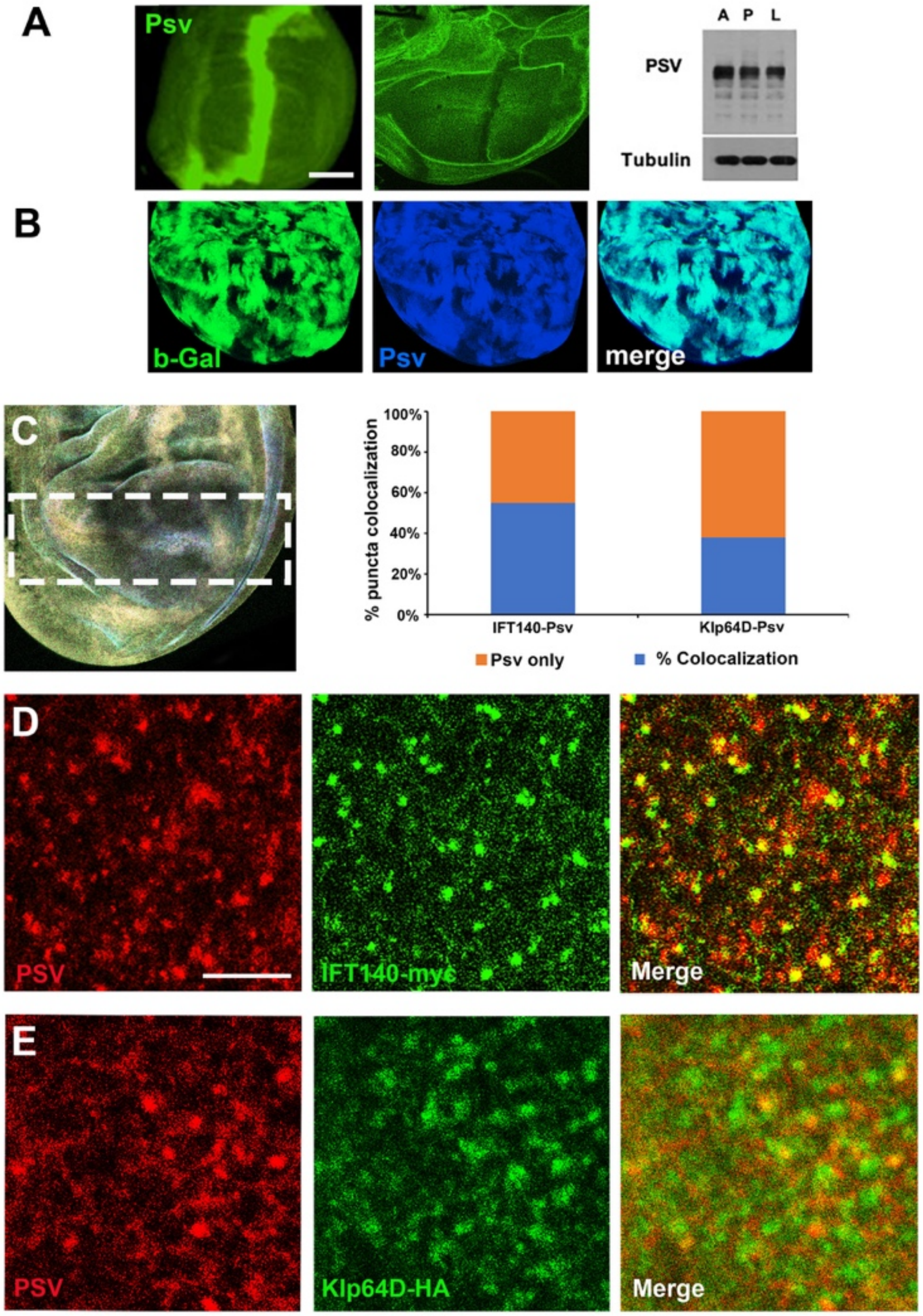


**Figure S4. Anti-Psv antibody characterization and overlapping localization of Psv with IFT140 and Kinesin2.**

**(A-B)** Generation and validation of an anti-Psv antiserum. **(A)** Left panel: Psv was detected at increased levels in the *patched* (*ptc*)-expressing stripe (*ptc-Gal4*) of the wing disc (*ptc>UAS-Psv*); note stripe of Psv expression. Middle panel: Conversely, Psv antibody showed a decreased level of Psv protein when *psv* was knocked down in the *ptc*-stripe (*ptc>psv RNAi)*. Right panel: Western blot analysis of protein extracted from adult flies (A), pupal (P) and 3$^{rd}$ larval (L) stages of *hs-Gal4/UAS-Psv* stained with anti-Psv antibody. **(B)** To confirm the specificity of the antibody wing discs carrying *psv*-/- LOF clones (marked by absence of β-Gal staining, green) were stained with anti-Psv (blue). Note absence of Psv staining in *psv*-/- LOF tissue patches.

**(C-E)** Co-localization of Psv and IFT140 or Kinesin2 in wing imaginal discs cells. **(C)** left panel shows region near the wing margin that was analyzed, and right panel shows quantification of co-localization. Note significant overlap with both proteins, IFT140 and Kinesin2, with IFT140 displaying a stronger co-localization. Examples of overlapping localization of Psv with IFT140-myc **(D)** and Klp64D-myc **(E)** in cells near the DV boundary region of wing discs. Wing discs from *C96>IFT140-myc* or *Klp64D-HA* were stained for Psv (red) and IFT140-myc (green), or Klp64D-HA (green). Scale bars in **A** and **D** indicate 50μm and 20μm, respectively. See also main Fig. 4 for additional information.

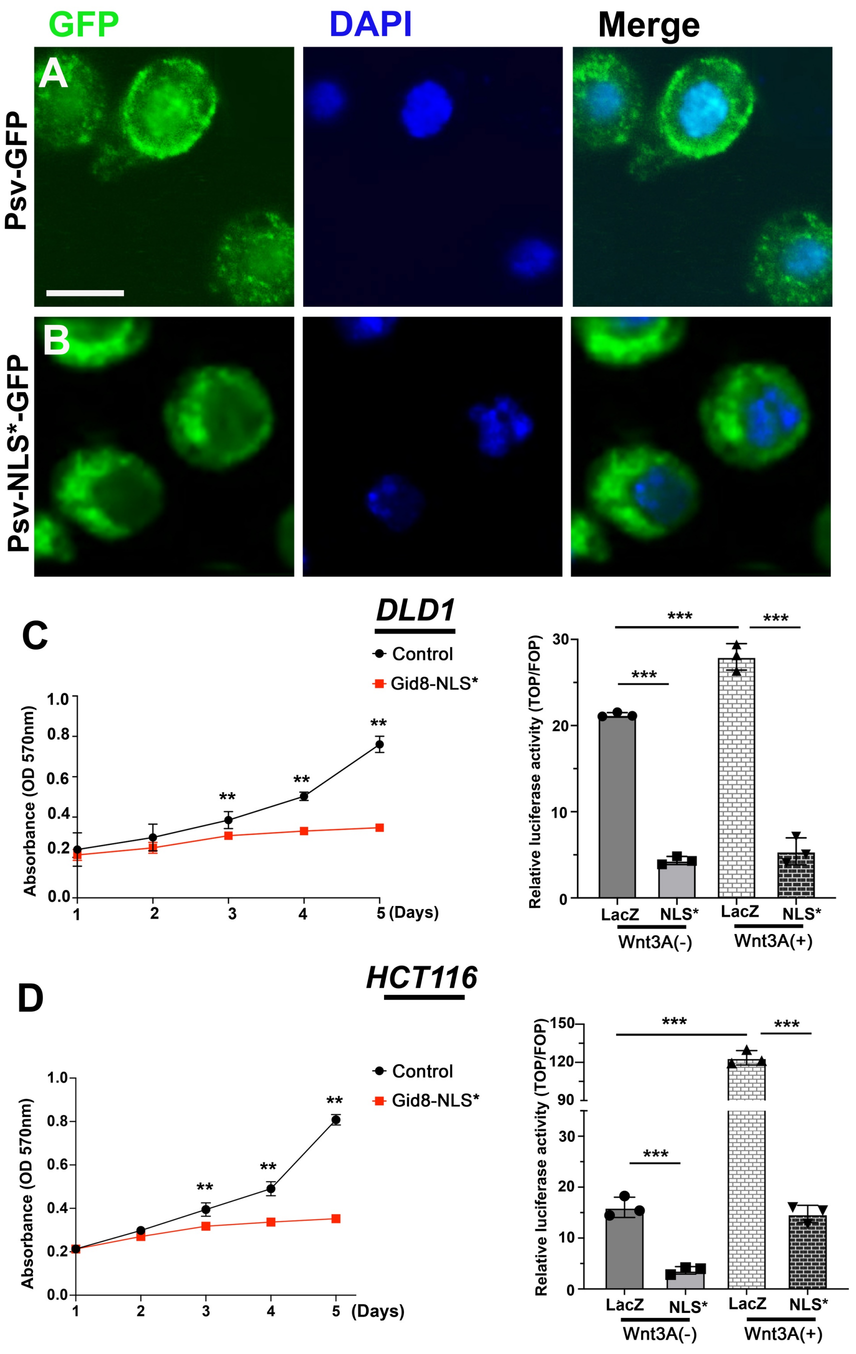


**Figure S5. Effects of PsvNLS mutation on Psv localization and function.**

**(A-B)** Transfection with Psv-GFP showed strong nuclear localization in S2 cells (**A**), whereas transfection of the Psv-NLS* mutation displayed largely cytoplasmic localization (**B**). Scale bar in **A** indicates 10 $\mu$m.

**(C-D)** Wnt-signaling in DLD1 cells (which carry a mutation in APC and thus are constitutively active) **(C)** and HCT116 (carrying a stabilizing mutation in $\beta$-catenin) **(D)** cancer cell lines (which both display constitutively active, ligand independent Wnt-signaling). Note that expression of the Gid8-NLS* mutant isoform markedly reduces growth of these cells (left graphs) and expression levels of the TOP/FOP Wnt signaling reporter (right graphs).

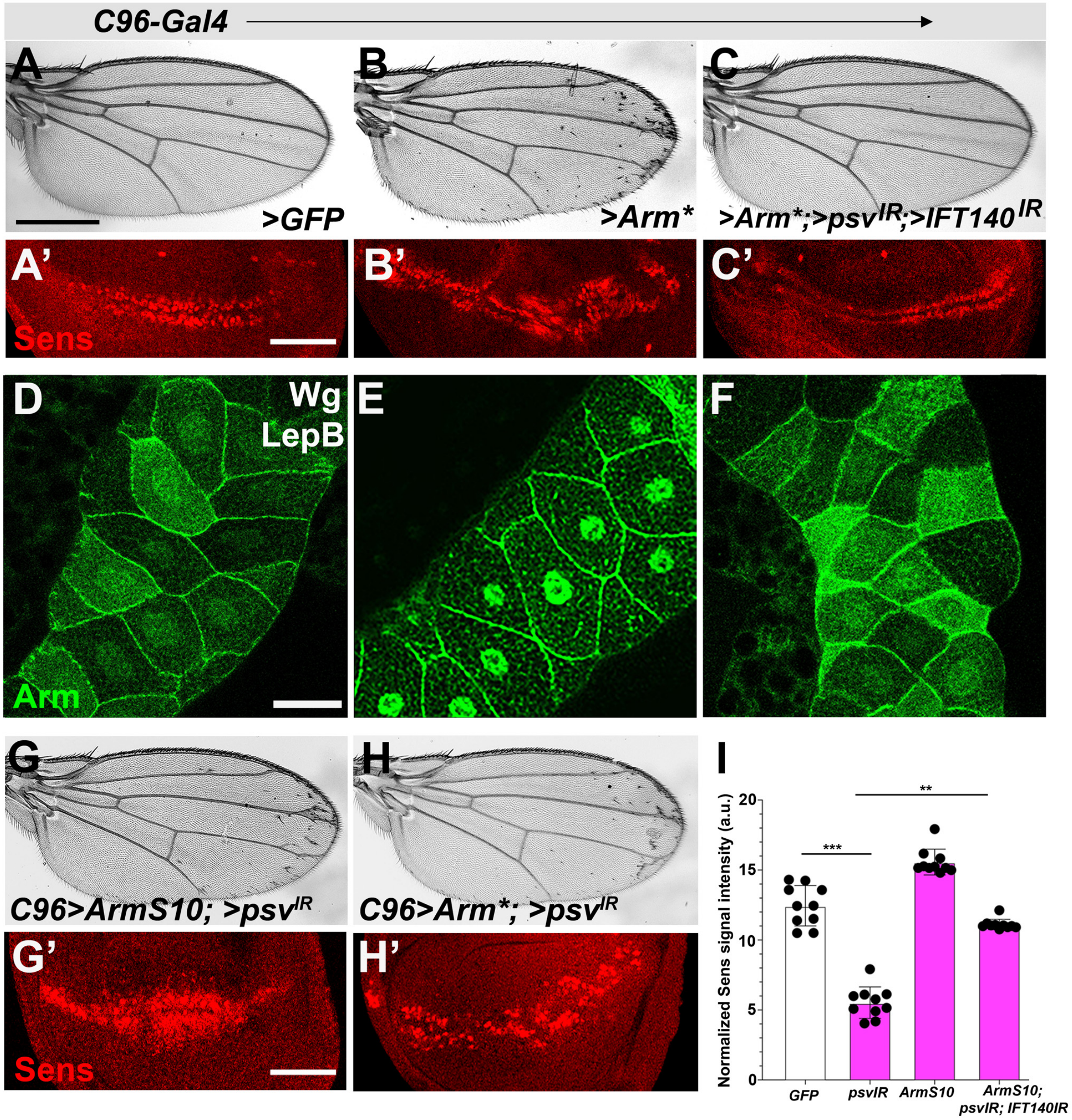


**Figure S6.** **Psv affects stabilized β-catenin mutations and inhibits Wnt signaling.**

**(A-A')** *UAS-GFP* (*C96>GFP*) control wing (A) and wing disc with wild-type Sens expression (red, **A'**) near D/V boundary. (**B-B'**) *C96>Arm** (a mutation in the priming phosphorylation site targeted by the destruction complex). Note GOF phenotype with ectopic margin bristle phenotype (**B**) and supernumerary ectopic Sens expressing cells in wing disc (**B'**). **(C-C')** *psv*$^{IR}$*; ift140*$^{IR}$ rescued the GOF phenotype caused by "activated, stable Arm" (Arm*).

**(D-F)** Assaying Arm* localization in the salivary glands (SGs, with Wg expression and LepB treatment). SGs were stained with anti-Arm, recognizing both endogenous Arm and Arm* (green). **(D)** Arm localizes to both membrane and nucleus (uneven Arm/β-catenin levels due to mosaic expression of Gal4 driver). **(E)** *C805>Arm**: shows increased Arm nuclear localization. **(F)** *C805>ArmS10, >psv*$^{IR}$*, >ift140*$^{IR}$: note markedly reduced Arm nuclear localization, although high levels of Arm staining (due to Arm* presence are detected in the cytoplasm (compare to panel **D**).

**(G-H')** Single knockdown of *psv* causes a partial rescue of the *C96>ArmS10* and *C96>Arm** phenotypes, as seen both in adult wings (**G, H**) and by Sens expression in wing discs (**G", H'**). Compare to figure panel **S6B** for Arm* and main Figure 6C, C' for ArmS10.

Scale bars in A-C and G-H indicate 700μm; A'-C' and G'-H' indicate 50 μm and D-F indicate 30 μm.

**(I)** Quantification of normalized Sens signal intensity, corresponding to panels in main Figure 6, as described in Suppl. Fig. S1.
